# Metabolic competition between lipid metabolism and histone methylation regulates sexual differentiation in human malaria parasites

**DOI:** 10.1101/2022.01.18.476397

**Authors:** Chantal T. Harris, Xinran Tong, Riward Campelo, Maria I. Marreiros, Leen N. Vanheer, Navid Nahiyaan, Vanessa A. Zuzarte-Luís, Kirk W. Deitsch, Maria M. Mota, Kyu Y. Rhee, Björn F.C. Kafsack

## Abstract

For *Plasmodium falciparum*, the most widespread and virulent malaria parasite that infects humans, persistence depends on continuous asexual replication in red blood cells, while transmission to their mosquito vector requires asexual blood-stage parasites to differentiate into non-replicating gametocytes. This decision is controlled by stochastic de-repression of a heterochromatin-silenced locus encoding *Pf*AP2-G, the master transcription factor of sexual differentiation. The frequency of *pfap2-g* de-repression was shown to be responsive to extracellular phospholipid precursors but the mechanism linking these metabolites to epigenetic regulation of *pfap2-g* was unknown. Here we show that this response is mediated by metabolic competition for the methyl donor S-adenosylmethionine between histone methyltransferases and phosphoethanolamine methyltransferase, a critical enzyme in the parasite’s pathway for *de novo* phosphatidylcholine synthesis. When phosphatidylcholine precursors are scarce, increased consumption of SAM for de novo phosphatidylcholine synthesis impairs maintenance of the histone methylation responsible for silencing *pfap2-g*, increasing the frequency of derepression and sexual differentiation.

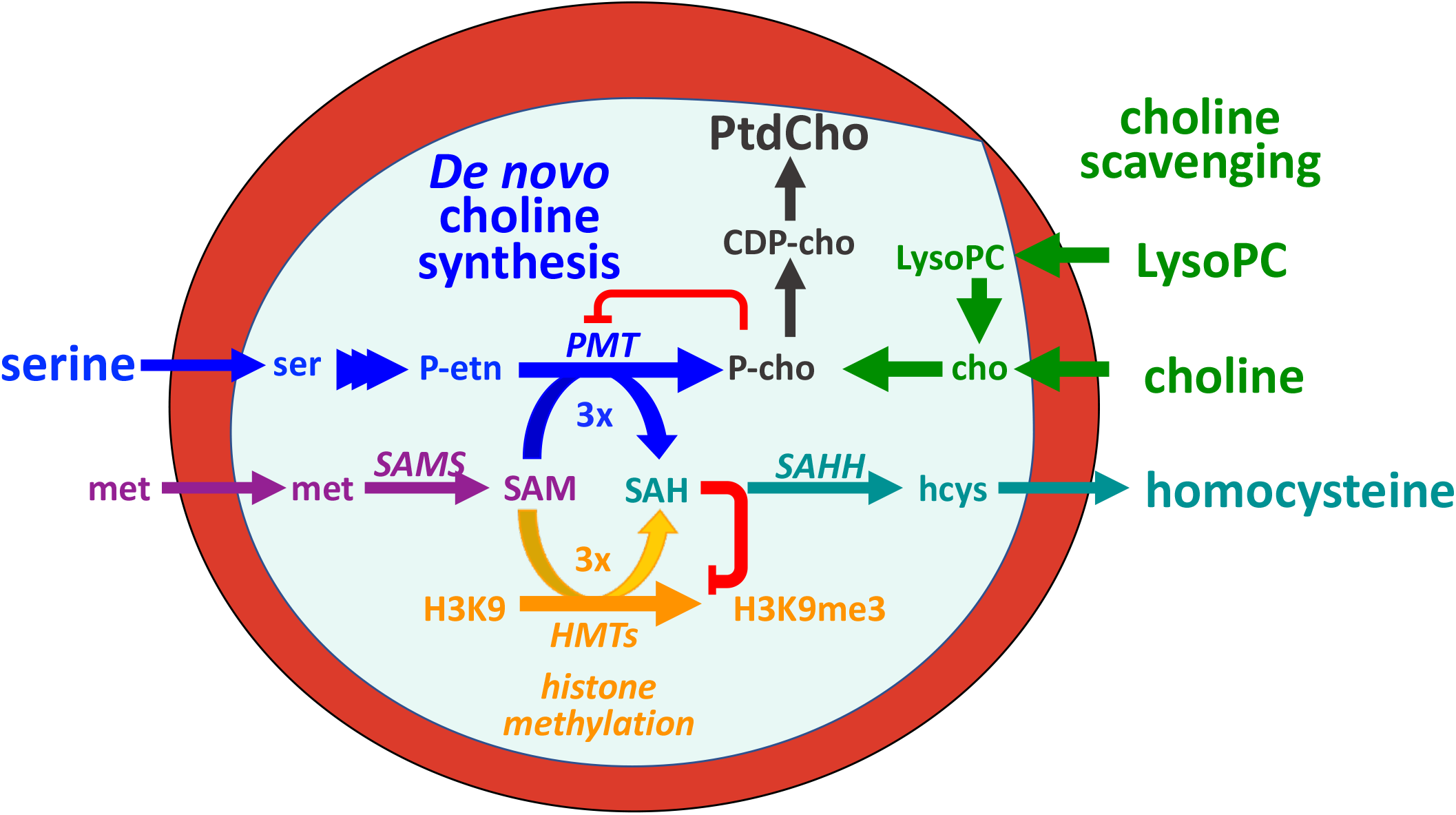

## INTRODUCTION

Transmission of malaria parasites requires differentiation from replicating asexual blood-stream forms that are responsible for pathogenesis into non-replicating sexual stages, called gametocytes, that can infect the mosquito vector. Parasite success depends on a careful balance between asexual replication, to maintain human infection, and gametocyte formation, to infect the mosquito vector for further transmission. The regulation of this balance is key to understanding the dynamics of malaria pathogenesis and transmission ^1^. Developmental commitment to sexual differentiation requires expression of the transcription factor AP2-G ^2,3^. This requires the transcriptional activation of the *ap2-g* locus, which is actively silenced in a heterochromatin-dependent manner during asexual replication ^2,4–6^. The frequency of sexual differentiation relies on effective heterochromatin maintenance at the *ap2-g* locus, which depends on the efficient trimethylation of lysine 9 on histone H3 of newly placed nucleosomes and recognition of this mark by heterochromatin protein 1 ^2,7–9^. So long as *ap2-g* remains efficiently silenced, parasites continue replicating asexually. However, AP2-G has several binding-sites within its own promoter that creates a positive transcriptional feedback loop once AP2-G expression rise beyond a threshold. When heterochromatin maintenance is impaired, leaky silencing allows AP2-G expression to exceed that threshold in a greater proportion of cells, leading to the activation of the feedback loop. This drives AP2-G expression to high levels, resulting in sexual commitment and activating the transcriptional program underlying gametocytogenesis ^2,10–13^.

In the laboratory, the frequency of sexual commitment of *Plasmodium falciparum*, the most wide-spread and virulent human malaria parasite, varies from less than 1% to greater than 40% in response to culture conditions, including parasite density and media composition ^14^. Recent work showed that lysophosphatidylcholine (LysoPC) and choline, precursors of phosphocholine (P-cho), the headgroup required for phosphatidylcholine (PtdCho) synthesis, act as potent suppressors of sexual commitment ^15^. This ability of parasites to vary their investment into gametocyte production illustrates the parasites’ ability to sense their in-host environment and respond adaptively to resource abundance ^16^ or anatomical niche ^17,18^. However, the underlying mechanisms that link the availability of these metabolites to *pfap2-g* activation remain unknown. Here, we show that the availability of P-cho precursors regulates sexual commitment by shifting the metabolic competition for the methyl donor S-adenosyl methionine (SAM) between the methyltransferase required for *de novo* synthesis of P-cho and the histone methyltransferases maintaining heterochromatin-mediated silencing, including at the *ap2-g* locus.

### P-cho precursor availability alters sexual commitment and intracellular concentrations of SAM and SAH

As the parasite grows during its 48-hour asexual replication cycle, the membrane content of infected red blood cells (iRBCs) increases 8-fold, with PtdCho accounting for more than half of this increase in membrane biomass ^19,20^. In *P. falciparum*, PtdCho is exclusively derived from P-cho, which can be generated *de novo* from serine via phosphoethanolamine (P-etn) or scavenged from extracellular sources, including choline and choline-containing phospholipids such as LysoPC ^21^ (Figure 1A, Extended Data Figure 1).

**Figure 1.**
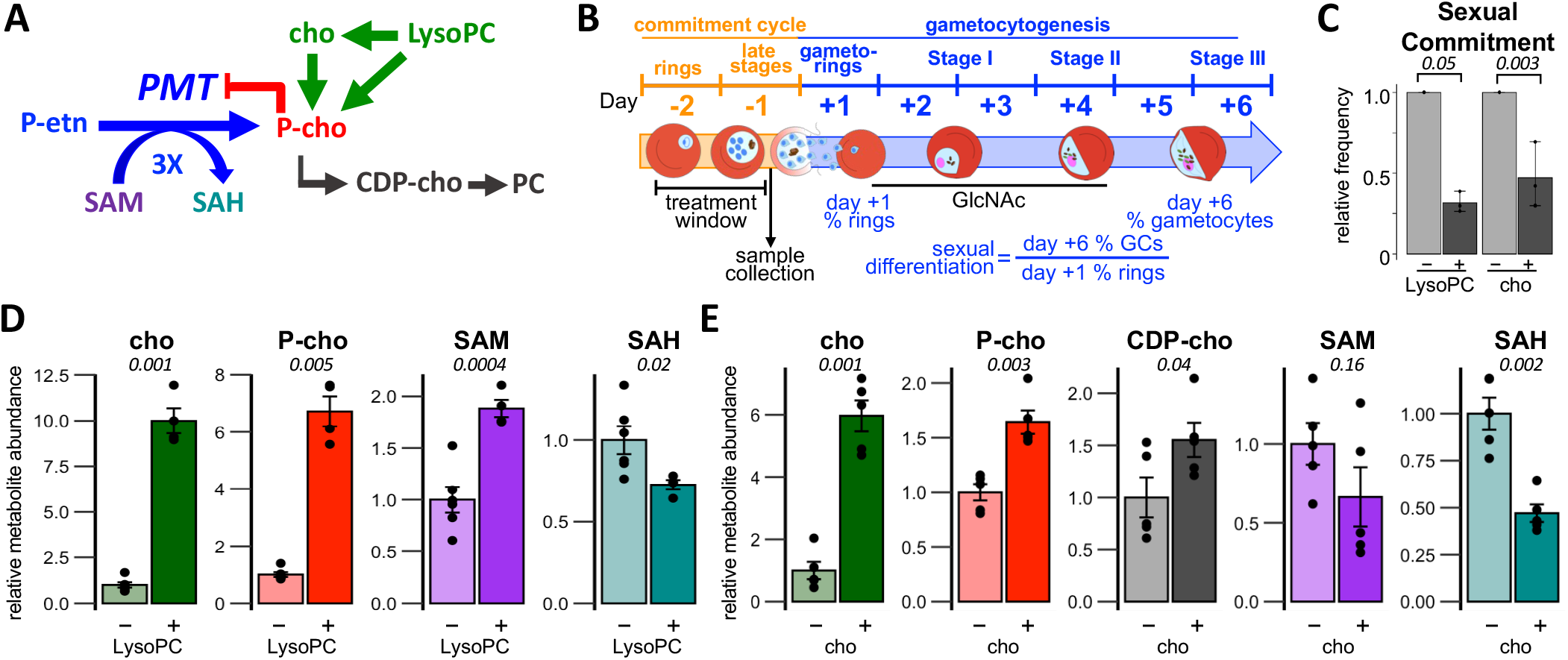
Phosphocholine precursor availability alters parasite SAM and SAH levels. **(A)** PtdCho (PC) is generated exclusively from P-cho, which can be scavenged from extracellular choline or LysoPC or synthesized de novo via triple methylation of P-etn by PMT, consuming 3 equivalents of SAM and producing 3 equivalents of SAH per P-cho. **(B)** Synchronous blood-stages were grown for a single cycle (commitment cycle) under various nutrient conditions and samples were collected 36 hours postinvasion. Treated parasites were allowed to re-invade and 50 mM N-acetylglucosamine was added on Day +1 to block asexual replication. The sexual differentiation rate is defined as the percentage of Day +1 ring stages that differentiate into stage III gametocytes by Day +6. **(C)** P-cho precursors inhibit the frequency of sexual differentiation (n=3). Italicized numbers are p-values from two-sided paired t-tests. **(D-E)** Intracellular metabolite levels are altered in response to LysoPC or choline supplementation in parasite media. (n=4-6). Bar graphs indicate the mean values relative to the reference condition ± s.e.m. Italicized numbers are p-values from two-sided t-tests.

Consistent with earlier reports ^15^, we found that supplementation of standard malaria growth media with exogenous P-cho precursors (see Extended Data Table 1 for media composition) during a single replication cycle substantially suppressed sexual commitment (Figure 1B-C) and increased the levels of P-cho and CDP-choline within iRBCs compared to conditions when these precursors were scarce (Figure 1D-E, Extended Data Figure 2). As we had previously described ^2^, the change in *ap2-g* expression at the end of this cycle tracked closely with the observed changes in sexual commitment (Extended Data Figure 3).

When exogenous P-cho precursors like choline or LysoPC are available for scavenging, parasite levels of SAH decreased significantly compared to conditions requiring greater *de novo* synthesis of P-cho from P-etn (Figure 1D-E). SAM levels increased in response to LysoPC but remained unchanged upon choline supplementation. A possible explanation for this difference is the additional energetic cost associated with scavenging choline, which must be phosphorylated using ATP, while LysoPC can be cleaved directly into P-cho by phospholipase C. This is supported by the observation that the suppressive effects of choline supplementation can be reversed by reducing the availability of glucose within the growth medium ^15^. This metabolic response to P-cho precursors is dose dependent (Extended Data Figure 2) indicating that P-cho precursor availability regulates intracellular levels of SAM and SAH. These changes in intracellular levels of P-cho, SAM, and SAH were limited to iRBCs and did not occur in uninfected RBCs (Extended Data Figure 4).

### Phosphoethanolamine methyltransferase activity drives the changes in intracellular SAM and SAH

Earlier work by the group of Choukri ben Mamoun showed that in malaria parasites *de novo* synthesis of PtdCho’s choline headgroup is carried out by (PMT), which generates P-cho by transferring three consecutive methyl-groups from the methyl-donor S-adenosylmethionine (SAM) onto P-etn ^21,22^, and that malaria blood stages substantially down-regulate *pmt* in response to supplementation with P-cho precursors ^23^ (Figure 2A). Since de novo synthesis of the choline headgroup is a major consumer of SAM and source of S-adenosyl homocysteine (SAH) in other eukaryotes ^24^, we wanted to assess whether the observed changes in SAM and SAH were due to a decrease in PMT activity. Consistent with the earlier reports, we observed a significant decrease in the abundance of the mono- and di-methylated P-etn reaction intermediates generated by PMT when cultures were supplemented with choline and found that PMT transcript abundance was substantially lower under this condition (Figure 2B). The effect of PMT activity on overall intracellular SAM and SAH levels was unclear but if P-cho synthesis by PMT is a major metabolic sink of SAM and source of SAH, their levels should correlate with PMT activity. To test this connection between PMT activity, SAM metabolism and sexual commitment, we generated inducible PMT knockdown parasites by inserting the glucosamine-responsive autocatalytic *glmS* ribozyme ^25^ into the endogenous PMT 3’UTR (Extended Data Figure 5). Treatment of PMT-*glmS* cultures with glucosamine during the commitment cycle reduced PMT protein levels by 55% (Figure 2D), resulting in a 2-fold increase in intracellular SAM and significantly suppressed sexual commitment, even in the absence of P-cho precursors (Figure 2E-F). This fits with earlier reports that found a near complete suppression of gametocytogenesis in *pfpmt* knockout parasites that was restored to parental levels upon genetic complementation ^25,26^.

**Figure 2.**
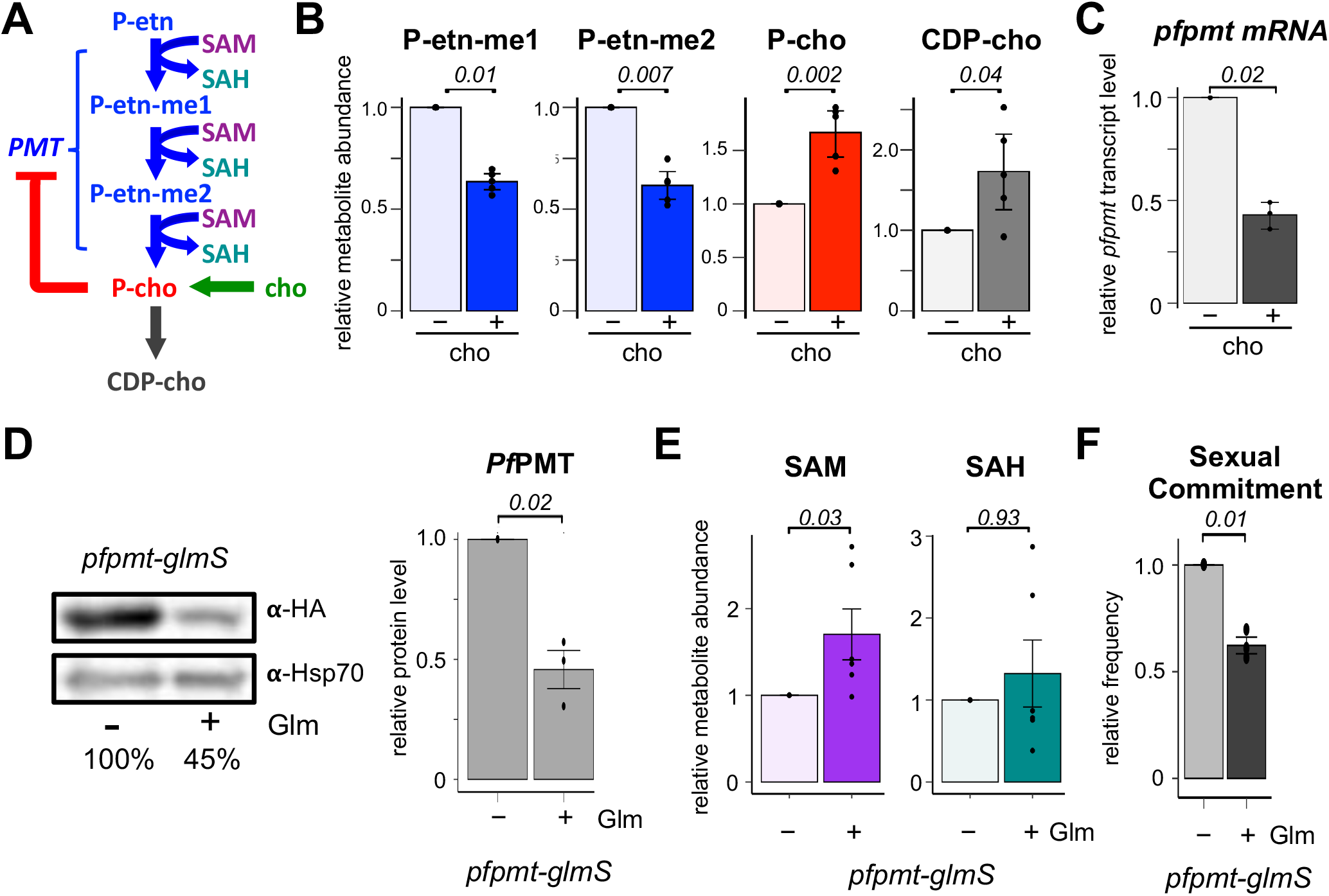
De novo synthesis of P-cho by *PMT* is a major sink of SAM and source of SAH. **(A-C)** Increases in P-choline from choline scavenging downregulates *PMT* activity as indicated by decreases in the monomethyl- and dimethyl-phopho-ethanolamine (P-etn-me1/2) reaction intermediates (**B**, n=4) and *PMT* transcript levels (**C**, n=3). Transcript abundance was normalized to seryl-tRNA synthetase transcript and shown relative to no choline supplementation. **(D)** Inducible knockdown of *PMT* upon addition of 2.5 mM glucosamine (Glm) in *PMT*-glmS parasites significantly reduces *PMT* protein abundance relative to *Pf*HSP70 loading control. (n=3) **(E)** Knockdown of *PMT* increases intracellular SAM levels (n=6). **(F)** Knockdown of *PMT* reduces sexual commitment even in the absence of P-cho precursors. (n=3) Bar graphs indicate the mean values relative to the reference condition ± s.e.m. Italicized numbers are p-values from two-sided paired t-tests.

### Intracellular SAM levels regulate sexual commitment

To determine whether these changes in SAM and SAH abundance merely correlate or directly regulate sexual commitment, we aimed to manipulate the intracellular concentrations of SAM and SAH independently of *PMT* activity and P-cho precursor abundance. We reasoned that if the availability of P-cho precursors suppresses sexual commitment by reducing *PMΓs* consumption of SAM, then this suppression should depend on the overall availability of SAM, which in malaria parasites is solely generated from methionine by SAM synthetase (SAMS). Malaria parasites are methionine auxotrophs and, unlike most eukaryotes, lack orthologs to enzymes able to regenerate methionine from homocysteine ^26^, making parasite SAM synthesis fully dependent on methionine derived from extracellular pools and the break-down of host cell proteins (Figure 3A). While removing extracellular methionine substantially decreased intracellular methionine and SAM (Figure 3B), it had no discernable effect on parasite replication over a two-week period (Extended Data Figure 6). However, while in standard growth medium containing 100 μM methionine, choline significantly suppressed commitment, this suppressive effect was greatly reduced when methionine was removed from the growth media for even a single commitment cycle (Figure 3C).

**Figure 3.**
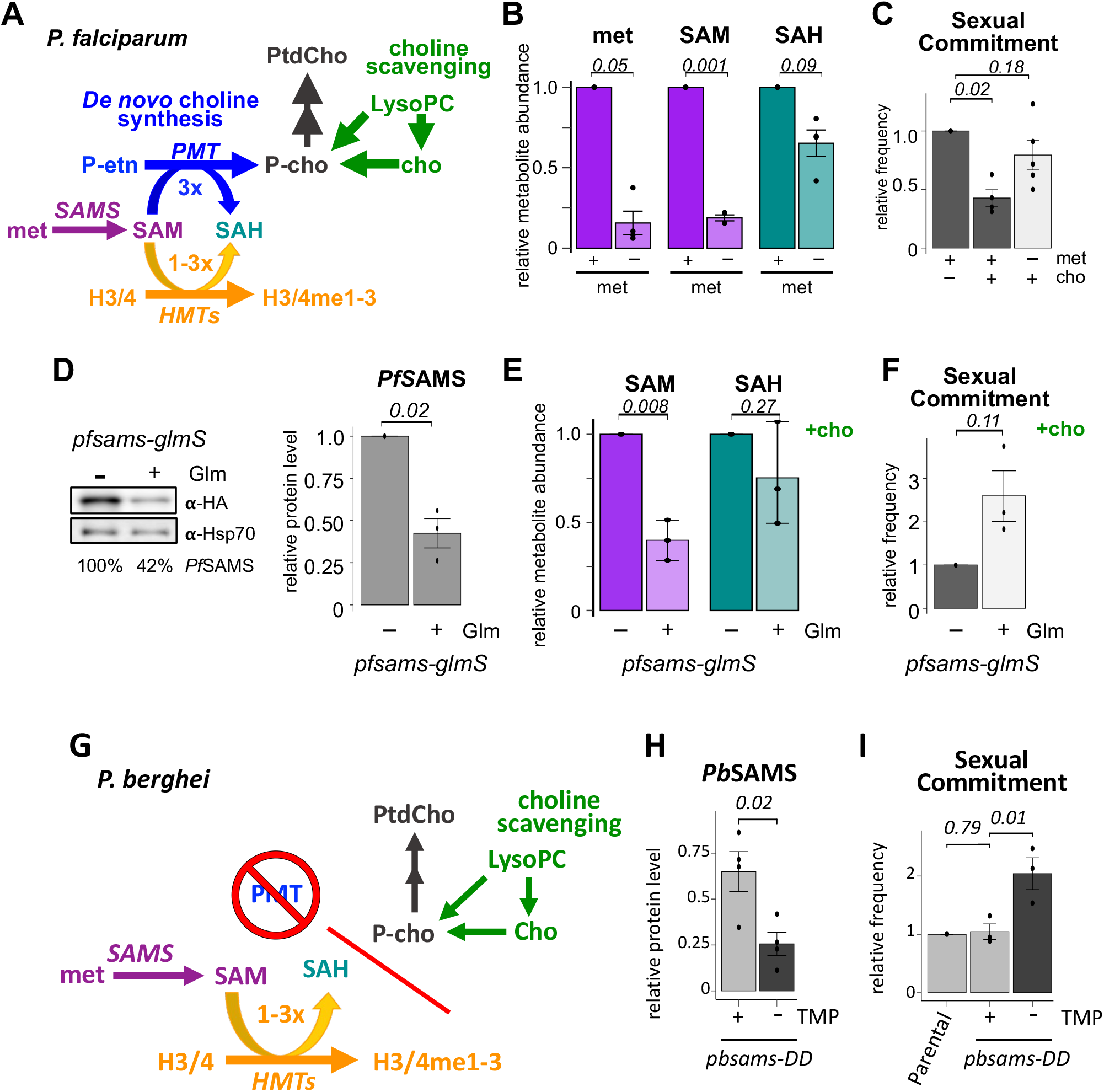
Intracellular SAM abundance regulates the frequency of sexual differentiation in human and rodent malaria parasites. **(A)** SAM and SAH are substrate and product, respectively, of both *de novo* P-cho synthesis and histone methylation. **(B)** Intracellular levels of methionine, SAM and SAH in growth medium containing standard (100 μM) or no methionine (n=3). **(C)** Removal of extracellular methionine reverses the suppressive effects of choline supplementation on sexual commitment (n=5). **(D)** Knockdown of *PfSAMS* in *pfsams-HA-glmS* parasites by treatment with 2.5 mM glucosamine (Glm) resulted in a 58% reduction in SAMS protein levels. **(E)** Knockdown of SAMS reduced intracellular SAM levels by 59% and **(F)** resulted in a 2.5-fold increase in sexual differentiation even in the presence of choline (n=3). **(G)** Loss of PMT in rodent malaria parasites, such as *P. berghei*, decouples SAM and SAH from PtdCho synthesis. **(H)** Removal of the stabilizing ligand TMP from *P. berghei pbsams-DD* knockdown parasites reduced *Pb*SAMS by 61% and **(I)** resulted in a 2-fold increase in sexual differentiation (n=3-4). Bar graphs show mean values relative to the reference condition ± s.e.m. Italicized numbers are p-values based on two-sided paired t-tests.

Since methionine is critical for a variety of cellular functions including translation, we wanted to test whether the effect of methionine on commitment levels was specific to changes in SAM availability. To this end we generated glucosamine-regulatable *pfsams-glmS* knockdown parasites (Extended Data Figure 7). Upon addition of glucosamine, protein levels of *Pf*SAMS were reduced by 58% (Figure 3D), which, even in the presence of choline, resulted in a two-fold reduction in SAM levels (Figure 3E) along with a more than 2.5-fold increase in sexual commitment (Figure 3F). Together, these findings demonstrate that sexual commitment is directly regulated by SAM availability.

### SAM, but not P-cho precursors, regulates sexual commitment in rodent malaria parasites

Earlier studies found that, unlike in *P. falciparum*, supplementation with P-cho precursors had little to no effect on sexual commitment in the rodent malaria parasite *Plasmodium berghei* ^15^. In the context of our proposed model, where the consumption of SAM by *PMT* provides the link between P-cho precursors and histone methylation, this observation is readily explained by the fact that the *PMT* ortholog was lost in the rodent malaria parasite lineage ^27^, thereby decoupling P-cho availability from SAM and SAH abundance in rodent parasites (Figure 3G). However, since heterochromatin-mediated silencing of the *ap2-g* locus also controls sexual commitment in rodent parasites ^3,5^, we hypothesized that commitment in *P. berghei* would never-the-less remain sensitive to changes in SAM availability. To test this, we generated SAMS knockdown parasites in *P. berghei* by creating a C-terminal fusion of the endogenous coding sequence with the ecDHFR-based destabilization domain (*pbsams-dd-ha*, Extended Data Figure 8A-B). Upon removal of the stabilizing ligand trimethoprim, *Pb*SAMS expression was reduced by 60% in *pbsams-dd-ha* blood-stages (Figure 3H, Extended Data Figure 8C) and resulted in a two-fold increase in sexual commitment (Figure 3I). This demonstrates that SAM availability regulates the rate of sexual commitment even when decoupled from PtdCho metabolism.

### Heterochromatin maintenance at the *pfap2-g* locus is responsive to changes in both SAM and SAH

In higher eukaryotes, changes in metabolism that alter intracellular SAM and SAH can have major effects on the methylation states of histone and associated regulation of gene expression ^24,28–31^. Methionine depletion and reduction in SAM abundance led to decreased methylation of specific histone modifications including H3K4me3, H3K9me3, H3K4me2, and H3K36me3, with H3K4me3 exhibiting the most profound changes, leading to a change in gene expression ^24,28,32^. Since the frequency of sexual commitment in malaria parasites is determined by the efficiency of heterochromatin-mediated silencing of AP2-G expression and changes in intracellular SAM directly alter commitment rates (Figure 3), we used CUT&RUN chromatin profiling ^33^ to examine whether these changes could substantially alter the abundance or distribution of the main histone methylation marks associated with both gene silencing (H3K9me3) and active transcription (H3K4me3) (Figure 4) ^34,35^. To evaluate the effect of SAM on histone methylation patterns we compared the distribution of these marks in the presence of abundant extracellular choline and methionine when SAM levels are high to conditions when the absence of these metabolites in the growth medium results in low SAM levels (Figure 4A), and elevated rates of sexual commitment (Figure 4B).

**Figure 4.**
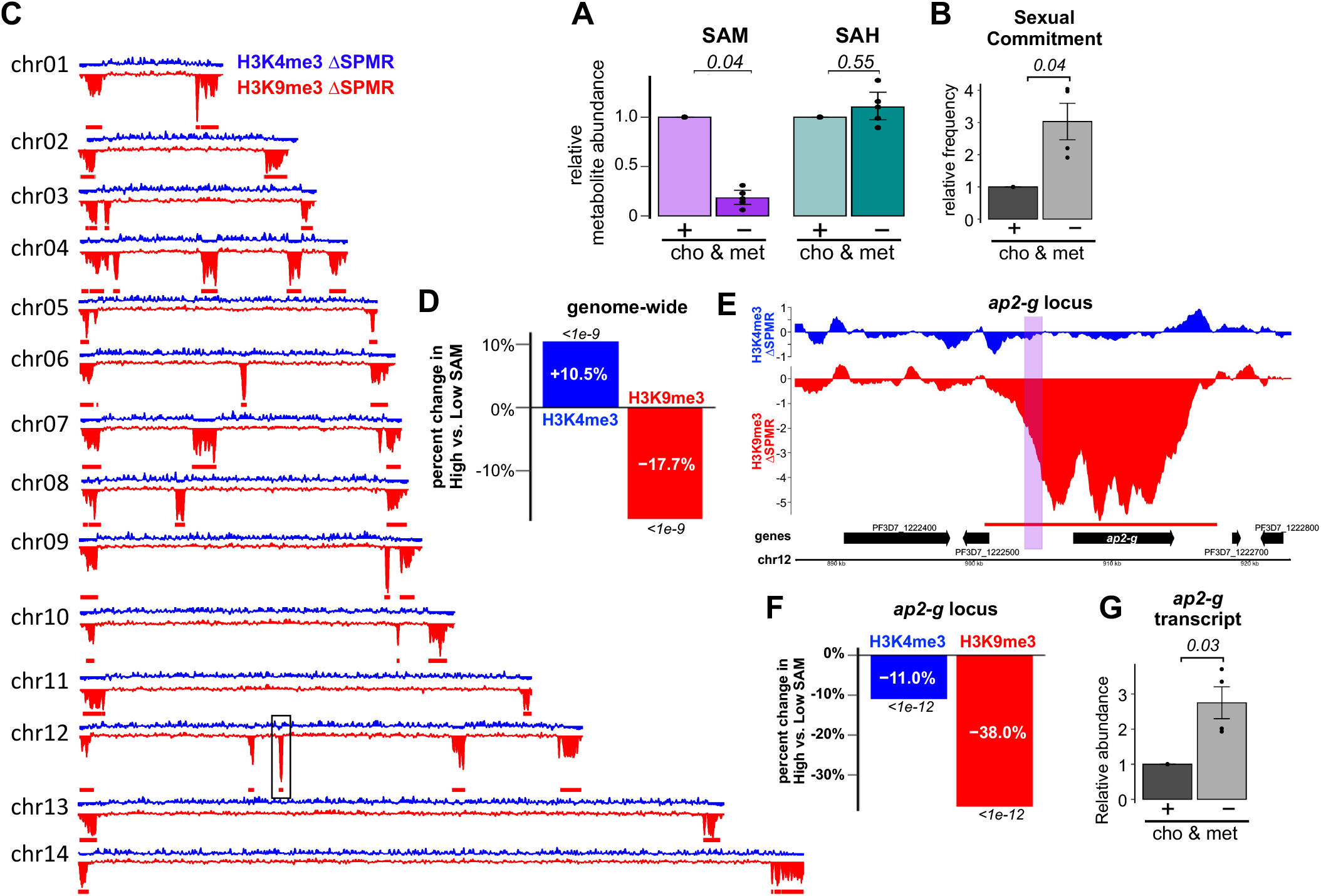
Reducing intracellular SAM levels impairs heterochromatin maintenance and increases both AP2-G expression and sexual commitment. **(A-B)** Relative intracellular abundances of SAM and SAH (n=5) **(A)**, and sexual commitment (n=4) **(B)** under high SAM (+cho & met) vs. low (-cho & met) SAM conditions. **(C)** Differences in H3K4me3 (blue) and H3K9me3 (red) abundance between parasites grown in low SAM versus high SAM conditions. The box on chromosome 12 indicates the *pfap2-g* locus. Red bars indicate regions of heterochromatin under high SAM conditions (n=2). SPMR: signal per million reads. **(D)** Genome-wide change in H3K4me3 (blue) and H3K9me3 (red) coverage under low SAM versus high SAM conditions. **(E)** Changes in the distribution of H3K4me3 (blue) and H3K9me3 (red) at the *pfap2-g* locus between parasites grown under low SAM versus high SAM conditions. The purple shaded region contains the AP2-G binding sites within the *ap2-g* promoter that drive the transcriptional feedback loop. **(F)** Change in coverage across the *pfap2-g* heterochromatin peak (red bar) of H3K4me3 (blue) and H3K9me3 (red) between parasites grown under low SAM versus high SAM conditions. **(G)** Relative abundance of *ap2-g* transcript levels between parasites grown under low SAM versus high SAM conditions (n=4).

In contrast to H3K4me3, which was somewhat elevated when intracellular SAM was low, we observed substantial genome-wide reductions in the abundance of the repressive H3K9me3 mark (Figure 4C-D) under low SAM conditions when compared to high SAM conditions. This reduction in H3K9me3 was observed in both regions of sub-telomeric heterochromatin domains and non-subtelomeric heterochromatin islands (Figure 4C, Extended Data Figure 9).

Closer examination of the region containing *pfap2-g* on chromosome 12 found that H3K9me3 occupancy across the locus was reduced by 38.0% under low SAM conditions (Figure 4E-F). At the leading edge of the heterochromatin island, which overlaps the promoter region containing the *Pf*AP2-G binding sites that drive the transcriptional feedback loop, this reduction was even more profound (−42.0% under low SAM, *p < 1e-12*). These reductions in the H3K9me3 silencing mark were accompanied by a 2.8-fold increase in transcript levels of *ap2-g* demonstrating that H3K9me3 occupancy and expression of the locus is responsive to changes in SAM (Figure 4G).

Each transfer of a methyl group from SAM generates one equivalent of SAH, which can act as a potent feedback inhibitor for many SAM-dependent methyltransferases ^36^, though histone methyltransferases differ in their susceptibility to feedback inhibition by SAH ^37^. Since *de novo* P-cho synthesis by *PMT* therefore represents not only a major sink of SAM but also an equivalently large source of intracellular SAH (Figure 2, Extended Data Figure 1), we also evaluated if direct changes to the intracellular levels of SAH can alter sexual commitment. To this end, we treated parasites during a single commitment cycle with 3-deaza-adenosine (3-DZA), a potent inhibitor of SAH hydrolase (SAHH), the enzyme that converts SAH into adenine and homocysteine (Figure 5A) ^38,39^. We found that treatment with 3-DZA increased commitment levels in a dose dependent manner while having only a minor effect on parasite growth (Extended Data Figure 10). Treatment with 100 nM 3-DZA in the presence of abundant methionine and choline greatly increased both SAH and SAM levels. The latter is likely the result of reduced consumption of SAM by methyltransferases as they are feedback inhibited by the increasing SAH levels (Figure 5B). SAHH inhibition by 3-DZA more than doubled sexual commitment, fully reversing the suppressive effects of choline supplementation (Figure 5C). These results demonstrate that elevating SAH levels is sufficient to increase sexual commitment rates, even when SAM is abundant. These data are in alignment with a previous study which found that homocysteine, the metabolic product and feedback inhibitor of SAHH that accumulates in culture and in the serum of infected patients, induces gametocytogenesis in culture ^40^.

**Figure 5.**
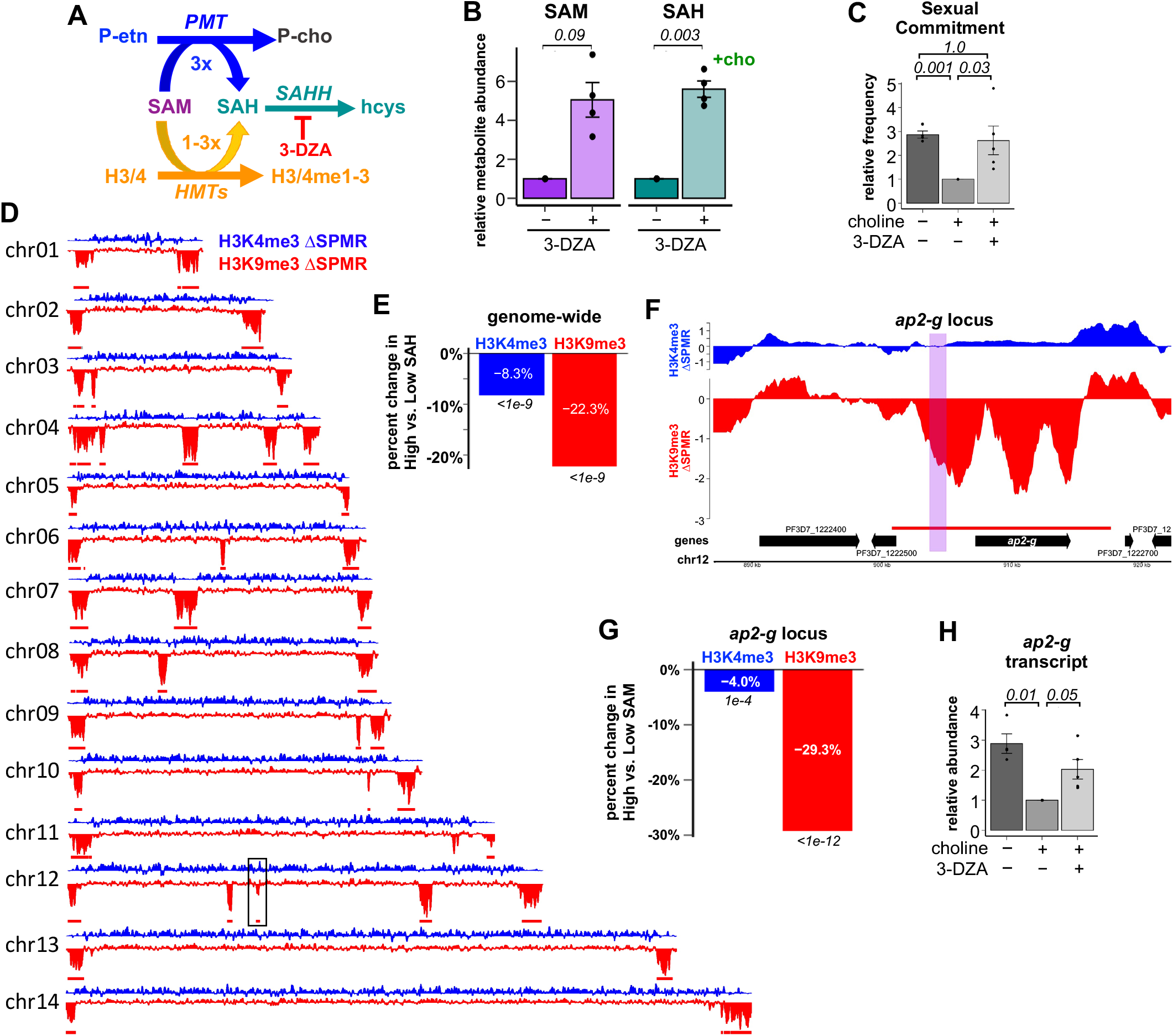
Increasing intracellular SAH impairs heterochromatin maintenance and increased both AP2-G expression and sexual commitment. Inhibition of SAHH with 3-DZA **(A)** increases intracellular SAM and SAH (n=5) **(B)** and sexual commitment (n=4) **(C)**. **(D)** Genome-wide differences in the distribution of H3K4me3 (blue) and H3K9me3 (red) between parasites grown in high SAH (+cho, +met, +3-DZA) versus low SAH (+cho, +met) condition as determined by CUT& RUN. The box on chromosome 12 indicates location of the *pfap2-g* locus. Red bars indicate regions of heterochromatin (n=2). SPMR: signal per million reads. **(E)** Mean genome-wide change in H3K4me3 (blue) and H3K9me3 (red) coverage in high SAH versus low SAH conditions. **(F)** Differences in the distribution of H3K4me3 (blue) and H3K9me3 (red) between parasites grown in high SAH versus low SAH conditions at the *pfap2-g* locus on chromosome 12. The purple shaded region contains key AP2-G binding sites that drive the transcriptional feedback loop. **(G)** Mean change in coverage across the *pfap2-g* heterochromatin peak (red bar) of H3K4me3 (blue) and H3K9me3 (red) between parasites grown in high SAH versus low SAH conditions. Italicized numbers are p-values based on two-sided paired t-tests for metabolite abundance or based on HOMER annotatePeaks and DESeq2 for histone modification abundance. **(H)** Relative abundance of *ap2-g* transcript levels between parasites grown under high SAH versus low SAH conditions (n=4).

Since suppression of commitment with choline reduced intracellular SAH levels but left SAM levels largely unchanged, we wanted to also evaluate whether high SAH was sufficient to impair heterochromatin maintenance even when SAM is abundantly available. While treatment with 3-DZA reduced genome-wide H3K4me3 abundance by 8.3%, it had even more profound effects on the H3K9me3 silencing mark, which was reduced 22.3% genome-wide (Figure 5D-E). This reduction in H3K9me3 was again observed in both regions of sub-telomeric heterochromatin domains and non-subtelomeric heterochromatin islands (Figure 4C, Extended Data Figure 11).

H3K9me3 occupancy across the *pfap2-g* locus on chromosome 12 was reduced by 29.3% under high SAH conditions (Figure 5F-G) and at the region containing the *Pf*AP2-G binding sites that drive the transcriptional feedback loop, this reduction was more profound at −39.2% (*p < 1e^-12^*) compared to low SAH conditions. These reductions in the H3K9me3 silencing mark were accompanied by a doubling of *ap2-g* transcript abundance (Figure 5H) demonstrating that H3K9me3 occupancy and expression of the locus are responsive to changes in SAH as well as SAM. Even when SAM is abundant, increasing SAH also impairs heterochromatin maintenance at the *pfap2-g* locus, increasing sexual commitment.

## DISCUSSION

Malaria parasites have evolved sophisticated mechanisms to sense and adapt to the diverse physiological niches they occupy during their life cycle. The switch from asexually replicating blood stage parasites to male and female gametocytes requires balancing a trade-off between maintaining in-host persistence and maximizing transmission between hosts. Several metabolites have been implicated in affecting this switch. Serine and homocysteine both increase frequencies of sexual differentiation ^19,40^, while LysoPC and its metabolic product, choline, both act as potent suppressors of sexual commitment ^15^. Additionally, the concentrations of these precursors are especially low in the bone marrow, the primary site of *P. falciparum* gametocytogenesis ^15,17^. Maturation in the bone marrow protects immature gametocytes from splenic clearance until they mature into highly deformable stage V gametocytes that are able to pass through the spleen ^41,42^. Intriguingly, increased sexual commitment of parasite infecting immature RBCs has been noted repeatedly ^43–45^. Unlike in mature RBCs, P-cho metabolism is highly active in immature RBCs and essential for terminal erythropoiesis ^46,47^. This would reduce the availability of P-cho precursors to parasites infecting immature RBCS and make them more reliant on *de novo* PtdCho synthesis via PMT.

Here we show that a drop in P-cho precursors leads to a decrease in intracellular SAM and rise in SAH as the result of increased *de novo* P-cho synthesis by PMT. These changes in SAM and SAH reduce the efficiency of H3K9me3 maintenance as the parasite replicates its genome 20-30 times during schizogony. This reduction in heterochromatin is not limited to the heterochromatin island on chromosome 12 that contains the *ap2-g* locus but occurs genome-wide. However, because AP2-G is the only transcription factor regulated by heterochromatin and because of its ability to bind the regulatory region of its own promoter, these reductions in heterochromatin can result in the activation of AP2-G and commit parasites to sexual development (Figure 6). However, because AP2-G is the only transcription factor regulated by heterochromatin and because of its ability to bind its own promoter, these reductions in heterochromatin can result in the activation of AP2-G and commit parasites to sexual development (Figure 6).

**Figure 6.**
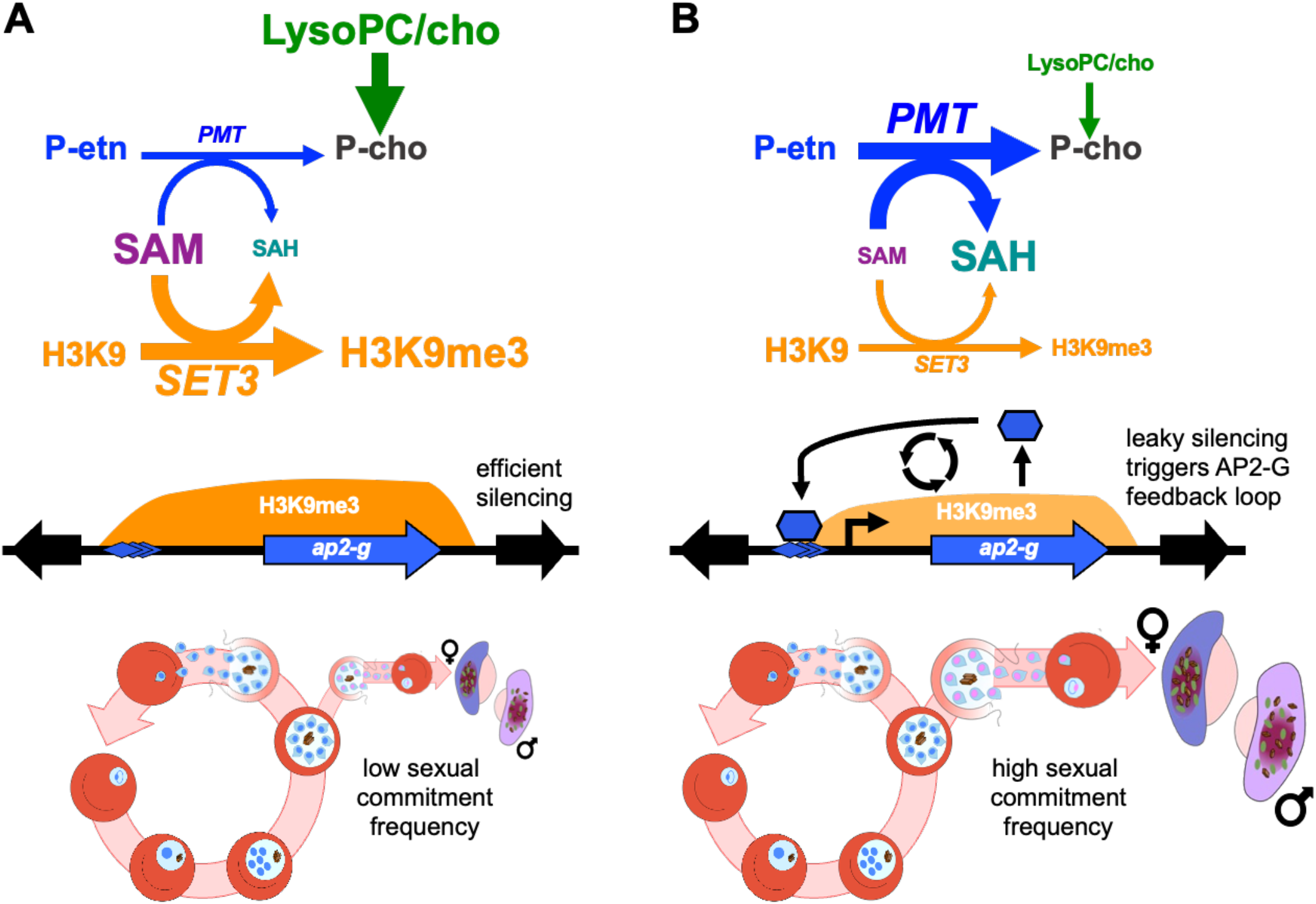
Metabolic competition between PMT and H3K9 methylation controls the rate of sexual commitment. **(A)** When P-cho precursors are available, H3K9me3 heterochromatin is efficiently maintained during schizogony resulting in low sexual commitment. **(B)** When P-cho precursors are scarce, increased *de novo P*-cho synthesis by *PMT* reduces SAM and increases SAH, both of which impair deposition H3K9me3 genome. Leaky silencing at the *pfap2-g* locus increases the probability of activating the positive transcriptional feedback loop, thereby increasing the frequency of commitment to sexual differentiation.

While we cannot exclude the possibility of other, unknown methylation reactions also regulating sexual commitment, H3K9 methylation has repeatedly been shown to strongly regulate AP2-G and sexual commitment in malaria parasites. Intriguingly, in malaria parasites methylation of H3K9 is more sensitive to changes in SAM and SAH than H3K4 methylation, unlike what has been reported for higher eukaryotes. This suggests a need for future examination of whether the K_m_ for SAM and K_i_ and SAH of *Pf*SET3, the parasite’s only Su(var)3-9 methyltransferase, are in the range of the intracellular concentrations of these two metabolites, which would allow their concentrations to directly regulate the deposition of the H3K9me3 silencing mark.

This model also explains the previously described observation that in rodent parasites sexual commitment is not responsive to P-cho precursors, as the loss of PMT in rodent malaria parasites decouples PtdCho synthesis from SAM, SAH, and histone methylation. Strikingly, this loss of PMT in rodent malaria parasites was accompanied by a lineage-specific expansion of the *fam-a* gene family from a single copy in the primate lineage to 42-215 copies ^48^. Members of this family contain the steroidogenic acute regulatory-related lipid transfer (START) domain and have been show to transport PtdCho ^49^. Without the ability to generate this key phospholipid *de novo*, this expansion presumably allows rodent malaria parasites to scavenge a full complement of PtdCho precursors.

In summary, changing availability of host LysoPC results in a shift in intracellular SAM/SAH which leads to changes in histone methylation, in particular the H3K9me3 silencing mark. Inefficient silencing at the *pfap2-g* locus increases the frequency of activation of the transcriptional feedback loop that commits parasites to sexual differentiation in *P. falciparum*. This allows parasites to increase sexual differentiation in the bone marrow where LysoPC is low and where gametocytes can develop without the risk of splenic clearance.

## ABBREVIATIONS

*3-DZA*: 3-deaza-adenosine
*cho*: choline
*CDP-cho*: cytidine diphosphate-choline
*CK*: choline kinase
*GlcNAC*: N-acetyl glucosamine
*Glm*: glucosamine
*hcys*: homocysteine
*iRBC*: infected *RBC*
*LysoPC*: lyso-phosphatidylcholine
*met*: methionine
*P-cho*: phospho-choline
*P-etn*: phospho-ethanolamine
*P-etn-me1/2*: mono/di-methylphosphoethanolamine
*PMT*: phosphoethanolamine methyltransferase
*PtdCho*: phosphatidylcholine
*RBC*: red blood cell
*SAH*: S-adenosylhomocysteine
*SAHH*: S-adenosylhomocysteine hydrolase
*SAM*: S-adenosylmethionine
*SAMS*: S-adenosylmethionine synthetase
*ser*: serine
*TMP*: trimethoprim
*uRBC*: uninfected RBC

## General

We wish to thank Dr. Jeffrey Dvorin (Boston Childrens Hospital) for providing Compound 1, the Weill Cornell Medicine genomics core for technical support, and the Eukaryotic Pathogen, Vector and Host Informatics Resource (VEuPathDB) for providing essential bioinformatics resources.

## Author contributions

Conceptualization: BFCK, KWD, MMM; Methodology: BFCK, CTH, MMM; Investigation: CTH, XT, RCM, LNV, IMM; Software, Formal Analysis, Data Curation: BFCK, CTH, XT; Writing – Original Draft: CTH Writing – Review & Editing: BFCK, CTH, MMM; Visualization: BFCK, CTH; Supervision: BFCK, KYR, MMM; Project Administration: B.F.C.K; Funding Acquisition: BFCK, KWD, MMM.

## Data and materials availability

All data needed to evaluate the conclusions in the paper are present in the paper or the supplementary materials. Raw and processed CUT & RUN data can be obtained from the NCBI Gene Expression Omnibus (GSE197916). The CUT & RUN analysis pipeline is available at https://github.com/KafsackLab/MetChoH3K9me3.

## MATERIALS AND METHODS

### *P. falciparum* cell culture

The parasite strains used for this study were NF54 obtained from BEI Resources and the *Pf-peg4-tdTomato* ^50^. Cultures were maintained using established culturing techniques ^51^. Standard complete media was RPMI-1640 supplemented with 25 mM HEPES, 368 μM hypoxanthine, 1 mM sodium hydroxide, 24 mM sodium bicarbonate, 21 μM gentamycin (Millipore Sigma), with Albumax II Lipid-Rich BSA (Gibco), unless otherwise stated. Concentrations for choline and LysoPC supplementation were used according to ^15^. See Extended Data Table 1 for changes to media compositions used throughout this manuscript. RBCs or cultures were washed three times with respective experimental media conditions at the start of each change in media condition.

### Generation of *Pf*SAMS-glmS and *PMT-glmS* conditional knockdown lines

The 3’ portion of the coding sequence for *pfsams* (577-1206bp) and *PMT* (651-1217bp) was amplified from gDNA and cloned by Gibson assembly into the pSLI-HAx3-glmS plasmid (gift from Professor R. Dzikowski) ^25^ following *NotI* and *XmaI* double digest. Following transfection of NF54 parasites, cultures were selected for the presence of the plasmid with 4nM WR99210 (gift of Jacobus Pharmaceuticals), follow by selection for integrants with G418 (Millipore Sigma). Single-crossover integration of the plasmid was confirmed via PCR (Extended Data Figures 5–6, Extended Data Table 2), and parasites were cloned to isolate a population with single cross-over lacking any remaining WT loci. Integrated parasite lines were maintained under G418 drug pressure.

### Animal Maintenance

Animal research was conducted at the Instituto de Medicina Molecular (Lisboa, Portugal). All protocols were approved by the animal ethics committee (ORBEA committee) at the institute and performed according to national and European regulations. BALB/c mice (age 5-8 weeks; males) were purchased from Charles River Laboratories (Saint-Germain-sur-l’Arbresle, France), kept in specific-pathogen-free conditions, and subjected to regular pathogen monitoring by sentinel screening. Experimental animals were randomly assigned and allowed free access to water and food.

### Generation of *Pb*SAMS-DD conditional knockdown parasites

Wild-type *P. berghei* ANKA strain was obtained from the MR4 repository (Manassas, Virginia). *P. berghei pbsams-DD* parasite line was obtained by double crossover homologous recombination. To do so, parasites were transfected by electroporation of purified schizonts, harvested on day 7-10 post transfection and genotyped by PCR. Transgenic parasites were then dilution cloned and further stored at −80 °C in frozen blood vials, containing 10^7^ blood stage parasites.

Recombinant parasites carrying the human dihydrofolate reductase (*hdhfr*) gene cassette were positively selected by treatment of mice with pyrimethamine and trimethoprim to stabilize *Pb*SAMS. Confirmation of transgenic parasite genotype, construct integration at the desired genomic loci and elimination of WT locus were assessed by PCR. Blood from the tail vein of infected mice was collected in 200μL of 1x PBS and genomic DNA was isolated using the NZY Blood gDNA Isolation Kit (NZYTech), according to manufacturer’s guidelines (Extended Data Figure 7, Extended Data Table 2). Stabilization of *Pb*SAMS-DD fusion protein throughout infection was achieved in vivo, by administration of trimethoprim (TMP) to mice (0.25 mg/ml of TMP in drinking water), 2 days prior to infection.

### Quantification of *Pb*SAMS knockdown

All steps of parasite pellet extraction protocol were performed at 4ºC to minimize protein degradation and all centrifugations were executed at 1000 x g for 10 min. Mice were sacrificed at day 4 post infection, 1mL of blood was collected by cardiac punction, washed in 10mL of 1x PBS and centrifuged. Packed erythrocytes were saponin-lysed in 0.15% saponin and centrifuged. Parasite pellet was washed twice in PBS containing 1x Proteinase inhibitor cocktail (Roche cOmplete Protease inhibitor tablets, EDTA free). Parasite pellet was then resuspended in lysis buffer (4 % SDS; 0.5 % Triton X-114 in 1x PBS), incubated on ice for 10 min, centrifuged at 21000 x g for 10 min and the supernatant was collected. Total protein content was determined using the Bio-Rad protein assay kit according to manufacturer’s instructions. Protein samples diluted in 5x SDS sample buffer (NZYTech) were denatured at 95º for 10 min and resolved in an 8% polyacrylamide gel (SDS-PAGE). Proteins were blotted into a nitrocellulose membrane by wet transfer at 200mA for 2 hours. Primary antibodies, mouse anti-HA antibody (1:1000, from Covance) and rabbit anti-Bip (1:2000, GeneScript) were incubated overnight at 4ºC. Secondary antibodies, anti-mouse horseradish peroxidase (HRP)-conjugated and anti-rabbit HRP (1:10000, Jackson ImmunoResearch Laboratories) were incubated at RT for 1 hour. Signal detection was obtained using Luminata Crescendo Western HRP substrate (Merck Milipore®) and the ChemiDoc XRS+ Gel Imaging System (BioRad). Protein band quantification was performed on Image Lab software (version 5.0) using BIP levels for normalization.

### Quantification of gametocytes in *Pb*SAMS-DD-HA parasites

Mice infections were performed by intraperitoneal inoculation of 1×10^6^ infected red blood cells (iRBCs) obtained by passage in the correspondent BALB/c background mice. Parasitemia (% of iRBCs) was monitored daily and gametocytemia (% of mature gametocytes) was determined on day 3 after infection, by microscopic analysis of Giemsa-stained blood smears. A total of 5-10 thousand RBCs were analysed in randomly acquired images and semi-automatically quantified using Image J software (http://rsbweb.nih.gov/ij).

### Induction and quantification of *P. falciparum* sexual commitment

Synchronous gametocyte induction was performed as previously described ^10^. Briefly, parasites were double-synchronized with 5% sorbitol to achieve a synchrony of ± 6h in the previous cycle. Synchronized ring-stage parasites were set up at 1.5% parasitemia (1% hematocrit) in 96-well, flat bottom plates under specific nutrient conditions to induce sexual commitment. Following reinvasion, on the first day of gametocyte development (D+1), ring-stage parasitemia was determined using flow cytometry and 50mM *N*-acetyl-D-glucosamine was added for 5 consecutive days. Gametocytes were then counted on day 6 (D+6) and commitment rate was determined by dividing the D+6 gametocytemia by the D+1 parasitemia assessed prior to *N*-acetyl-D-glucosamine addition. Gametocytes induced using the Pf-*peg4*-tdTomato fluorescent gametocyte reporter line were counted using flow cytometry ^50^.

### RNA Extraction, cDNA synthesis, and quantitative RT-PCR

Total RNA from saponin-lysed parasites was extracted using Trizol (Invitrogen) or Direct-Zol RNA MiniPrep Plus kit (Zymo Research). 500ng total RNA was pre-treated with 2U amplification grade DNase I (Invitrogen) and reverse transcribed using SuperScript III Reverse Transcriptase kit (Invitrogen) and random hexamers (Invitrogen). Quantitative PCR was performed on the Quant Studio 6 Flex (Thermo Fisher) using iTaq Sybr Green (Bio-Rad) with specific primers for selected target genes (Extended Data Table 2) and normalized to seryl-tRNA synthetase or ubiquitin-conjugating enzyme transcript abundance. To quantify AP2-G transcript levels at the very end of the commitment cycle, 2.5 μM Compound 1 (a generous gift of Dr. Jeffrey Dvorin) was added at 36 hpi to prevent egress and RNA was extracted 12 hours later.

### Metabolite Analysis

Synchronous *P. falciparum* parasites were grown from 8hpi to 34hpi ± 6 hpi (100mL, 3% hematocrit, 5-6% parasitemia) under the indicated growth conditions. Infected red blood cells (iRBC) were then purified to > 90% purity using a 70/40% percoll/sorbitol density gradient and centrifugation (4700 x g for 15 minutes at room temperature), then washed three times with minimal RPMI. Following isolation, equal numbers of isolated iRBC, as determined by the Beckman Coulter Z1 Coulter Particle Counter, were then re-incubated in their respective treatment conditions for four to six more hours to allow parasites to recover from percoll/sorbitol isolation. To ensure, that metabolites were extracted at similar points in the cell cycle, nuclear replication was followed by flow cytometry after staining with Hoechst 33342 (16 μM for 20 min) with a target nuclear content of 6-8N relative to a ring stage standard. Following recovery, the purity of iRBCs was assessed using flow cytometry, cells were then pelleted, washed once with 1 mL of 1x PBS, and quickly lysed with 500 μL of 90% ultrapure HPLC Grade methanol (VWR), followed by exactly 10 seconds of vortexing before being stored in dry ice. Lysed samples were then centrifuged at top speed at 4°C and supernatant collected for metabolite analysis. Uninfected red blood cells (uRBC) were incubated under the same treatment conditions as iRBC, counted, and methanol extracted to distinguish iRBC metabolite signal from uRBC signal. Extracted metabolites were stored at −80°C. LC-MS based metabolomic analysis was performed as previously described ^52^. 10 μL of extract was separated on an Agilent 1290 Infinity LC system containing a Cogent Diamond Hydride Type C silica column (150 mm × 2.1 mm; Microsolv Technologies). Acquisition was performed on an Agilent 6230 TOF mass spectrometer (Agilent Technologies, Santa Clara, CA) employing an Agilent Jet Stream electrospray ionization source (Agilent Technologies, Santa Clara, CA) operated in high resolution, positive mode. Metabolite identification was verified by exact mass (to 35 ppm) and co-elution with authentic standards purchased from Millipore Sigma. Batch feature extraction and chromatographic alignment across multiple data files was performed using the Agilent MassHunter Profinder software and extracted metabolite data was exported for further statistical analysis using R. Metabolites were quantified based on normalized peak area (peak area/(number of parasites extracted*dilution factor)).

We took extensive efforts to ensure that metabolite measurements were comparable. First, the nuclear content of all cultures was monitored closely by volumetric flow cytometry to ensure that cultures were at similar points in the cell cycle prior to extraction and to provide highly accurate measurement of the number of parasites extracted. Second, metabolite extraction was always carried out in small batched and samples to be compares were always extracted in parallel. Third, metabolite peak areas were normalized by the total of parasites extracted, dilution factor, and injection volume. Fourth, only samples included on the same LC-MS run were compared. Lastly, a shared reference sample with metabolite spike-ins was also included on each LC-MS run to facilitate comparison and account for differences in retention time.

### Western Blotting

*PMT*-glmS and *Pf*SAMS-glmS knockdown western blots were performed using whole cell lysate. Ring-stage parasites (12hpi ± 6 hpi) were treated with or without 2.5mM glucosamine (VWR) to induce protein knockdown. Parasites were saponin lysed at 36hpi ± 6 hpi and whole cell extract collected in SDS sample buffer, then boiled for 10 minutes. Proteins were then separated on 12% SDS PAGE and transferred to a PVDF membrane (Millipore Sigma, 0.2 μM). Membranes blocked with 5% milk were then probed with anti-HA (1:2000, Abcam, ab9110), and anti-hsp70 (1:2000, StressMarq SPtdCho-186) primary antibody solutions, followed by anti-rabbit IgG (1:5000, Millipore Sigma 12-348) secondary. Chemiluminescence was measured using the Azure c-Series imaging systems (Azure Biosystems) and quantified using the Fiji open-source image processing package based on ImageJ.

### Quantification of changes in H3K9me3 and H3K4me3 abundance and distribution

H3K9me3 and H3K4me3 abundances were measured by Cleavage Under Targets & Release Using Nuclease (CUT & RUN) ^53^ adapted for *P. falciparum* ^33^. Approximately 1 x 10^7^ percoll/sorbitol isolated schizonts (34-38 hpi) per sample were washed three times with 1 ml wash buffer (20 mM HEPES at pH 7.5, 150 mM NaCl, 0.5 mM Spermidine, and 1 × Roche complete protease inhibitor) at room temperature and resuspended in 225ul of wash buffer. 25ul of Concanavalin A-coated beads were washed and resuspended in binding buffer (20 mM HEPES–KOH at pH 7.5, 10 mM KCl, 1 mM CaCl2, 1 mM MnCl2) before being added to each sample. Samples were then rotated for 10 minutes at room temperature. Then, samples were placed on a magnetic stand to clear (30s to 2 minutes) and remove all the liquid. Samples were washed 3x’s with DIG wash buffer (wash buffer with 0.025% digitonin). For each sample, 150μl of antibody wash buffer was added (wash buffer, 0.025% digitonin, and 2 mM EDTA at pH 8.0) with H3K4me3 (0.005μg/μl, C15410003-50, Diagenode), H3K9me3 (0.005μg/μl, ab8898, abcam), or IgG isotype control (0.005μg/μl, 02-6102, ThermoFisher) was added to the sample tube and incubated at 4 °C, rotating, overnight. Following antibody incubation, samples were placed on a magnetic stand to clear and pull off the liquid, then washed 3 x’s with DIG wash buffer, 150 μl of ProteinA/G-MNASE fusion protein (dilution 1:60, 15-1016, EpiCypher) in DIG wash buffer and rotated at 4 °C for 1 hour. After washing samples 3x’s with DIG wash buffer, samples were then washed 3x’s with Low Salt Rinse Buffer (20 mM HEPES–NaOH at pH 7.5, 0.025% digitonin, 0.5 mM spermidine). 200 μl of ice-cold incubation buffer (3.5 mM HEPES–NaOH at pH 7.5, 100mM CaCl_2_, 0.025% DIG) was added and samples were equilibrated at 0 °C for 30 minutes to achieve targeted digestion. Digestion was then stopped by adding 200 μl of stop buffer (170 mM NaCl, 20 mM EGTA at pH 8.0, 20 mM EGTA, 0.0.25% digitonin, 25 μg/mL glycogen, 50 μg/mL RNase A). After incubation at 37 °C for 30 minutes to release soluble fragments, samples were digested by adding 2.5 μL proteinase K (20 mg/ml) and 2 μL 10% SDS, and then incubated at 50 °C for 1 hour. DNA fragments were purified with phenol–chloroform–isoamyl alcohol and washed by ethanol precipitation, and finally dissolved in 30 μL TE buffer (1mM Tris-HCl pH 8, 0.1 mM EDTA).

Library construction was carried out using NEBNext Ultra II DNA Library Prep Kit for Illumina (E7645S, NEB). 4ng of fragmented DNA was used for end-repair/A-tailing, ligation, and post ligation cleanup with 1.7x volumes of AMPure XP beads (Catalog number, company). Following cleanup, PCR amplification was performed using 2x KAPA HotStart ready mix (Catalog number, company) and NETFLEX primer mix (Catalog number, Bio Scientific) with PCR program: 1 min @ 98° C/15 cycles: 10 sec @ 98° C/1 min @ 65° C// 5min @ 65° C// hold 4° C. PCR products were size selected with 0.8x volumes, then 1.2x volumes of AMPure XP beads. Beads were then washed twice with 80% ethanol and DNA eluted with 0.1x TE and used for sequencing in a Nextseq2000 system as paired-end reads, following quality control.

After sequencing, raw reads were trimmed using Trimmomatic v0.38 ^54^ to remove residual adapter sequences and low quality leading and trailing bases. Both paired and unpaired reads were retained for read lengths that were at least 30 bases after trimming. Trimmed paired reads were aligned to the PlasmoDB version 46 *P. falciparum* 3D7 reference genome ^26^ using BWA v.0.7.1 ^55^. SAMtools v.1.10 ^56^ was used to remove low quality alignments as well as sort and index sample files. Normalized fold enrichment tracks were generated by using the MACS2 v.2.2.7.1 ^57^ callpeak function with settings: -f BAMPE -B -g 2.3e7 -q 0.05 – nomodel –broad –keep-dup auto –max gap 500. Bedgraph outputs were then passed into the bdgcmp function with the setting -m FE (fold enrichment) to generate signal tracks to profile histone modification enrichment levels compared to whole genome. Peak sets from replicates were compared with Bedtools intersect v2.26.0 ^58^, and peaks that overlapped by at least 1 bp were considered shared. Fold enrichment bedgraphs and peak sets were then output to RStudio Server (v1.4.1717) for further analysis using the GenomicRanges Package v.1.44.0 ^59^ and visualized with the GViz v1.38.1 package ^60^ within the Bioconductor project (release 3.13) ^61^. The full analysis pipeline can be found at https://github.com/KafsackLab/MetChoH3K9me3.

## EXTENDED DATA FIGURES

**Extended Data Figure 1:**
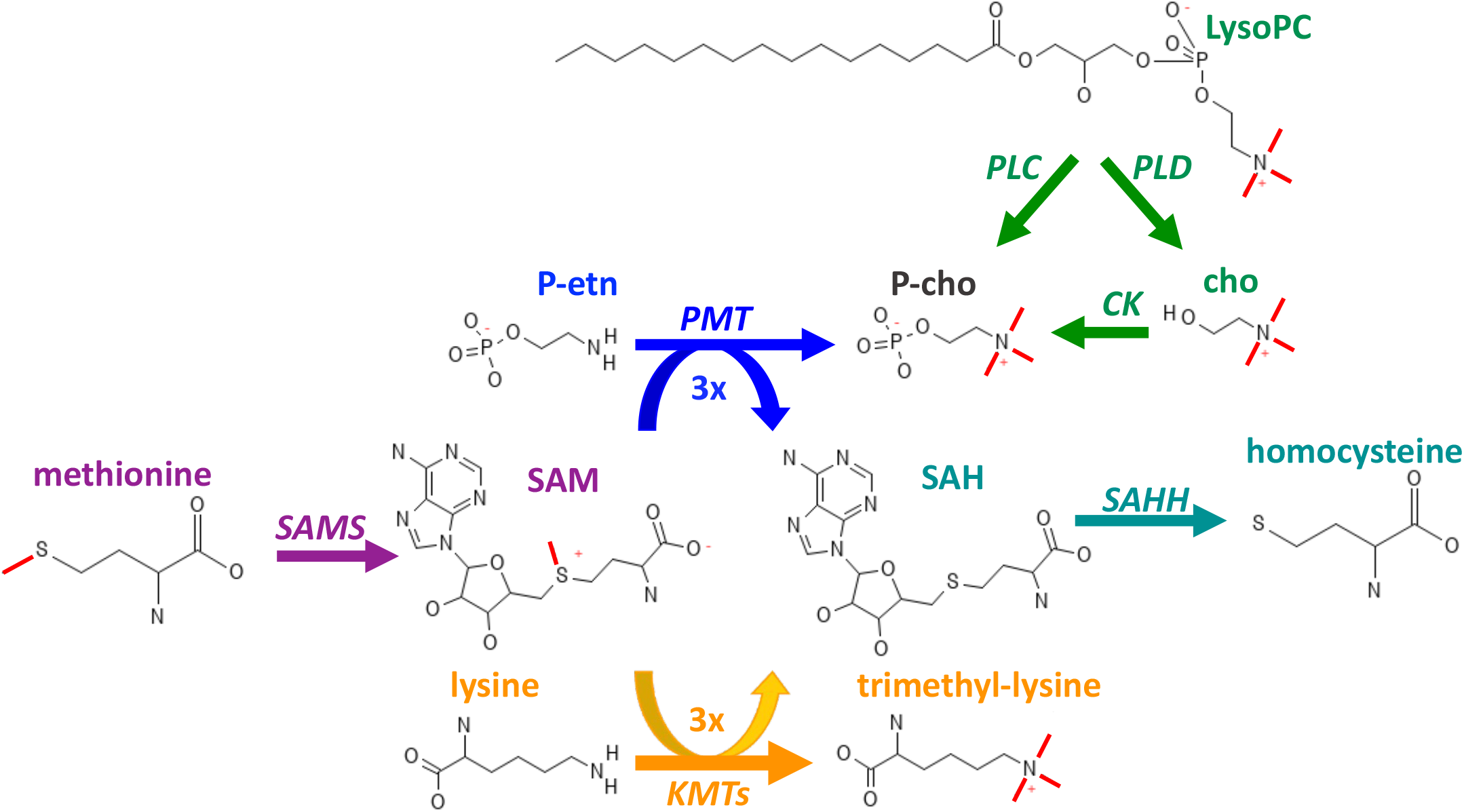
Central methylation-related metabolites in this study. Names of enzymes involved in their interconversion are noted in italics and methyl groups being transferred are highlighted in red.

**Extended Data Figure 2:**
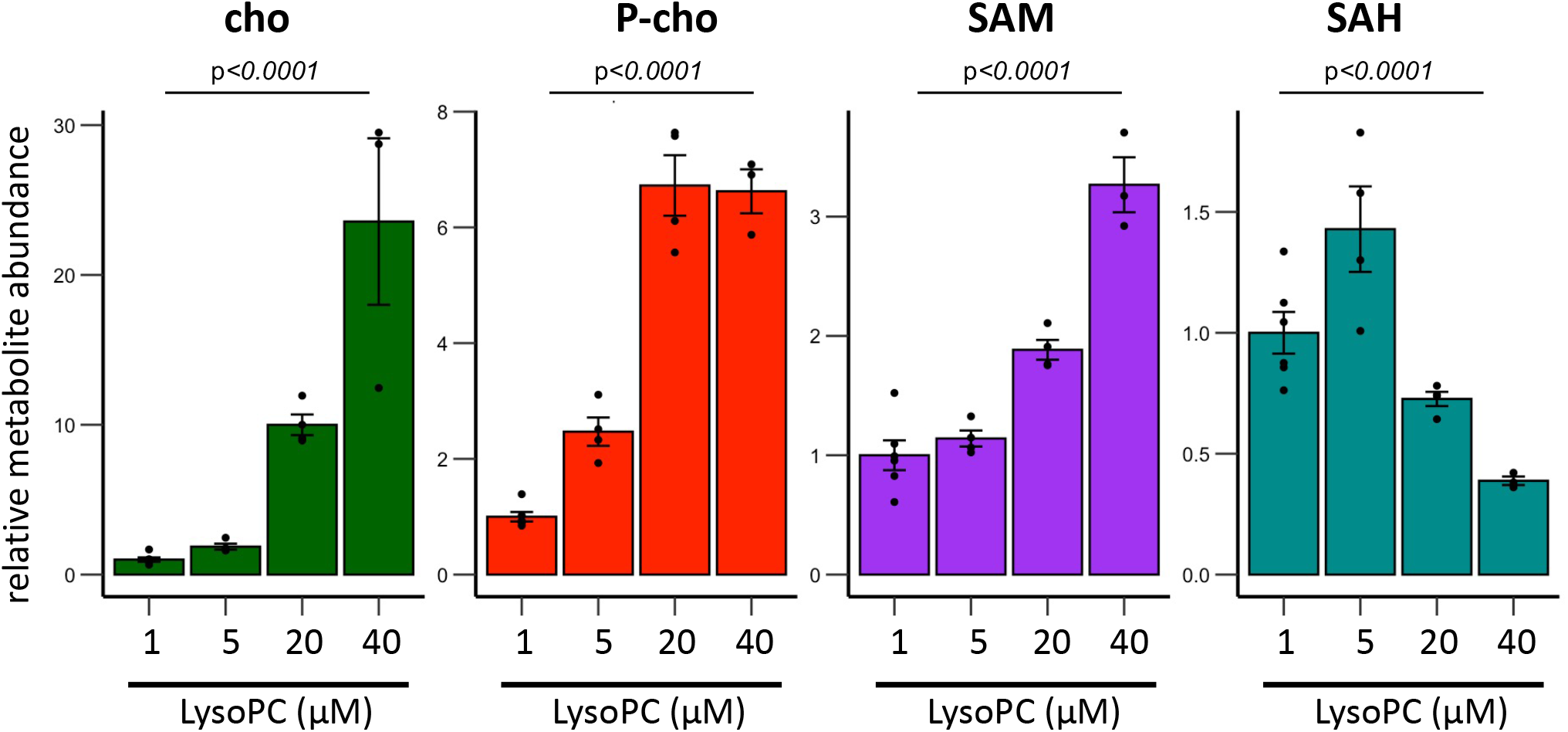
Dose-dependent metabolic response to LysoPC. Parasites were cultured in media spiked with increasing concentrations of LysoPC. Bar graphs show the mean intracellular metabolite abundances per thousand parasites ± s.e.m (n=3-5). Italicized numbers are p-values based on two-sided ANOVA tests.

**Extended Data Figure 3.**
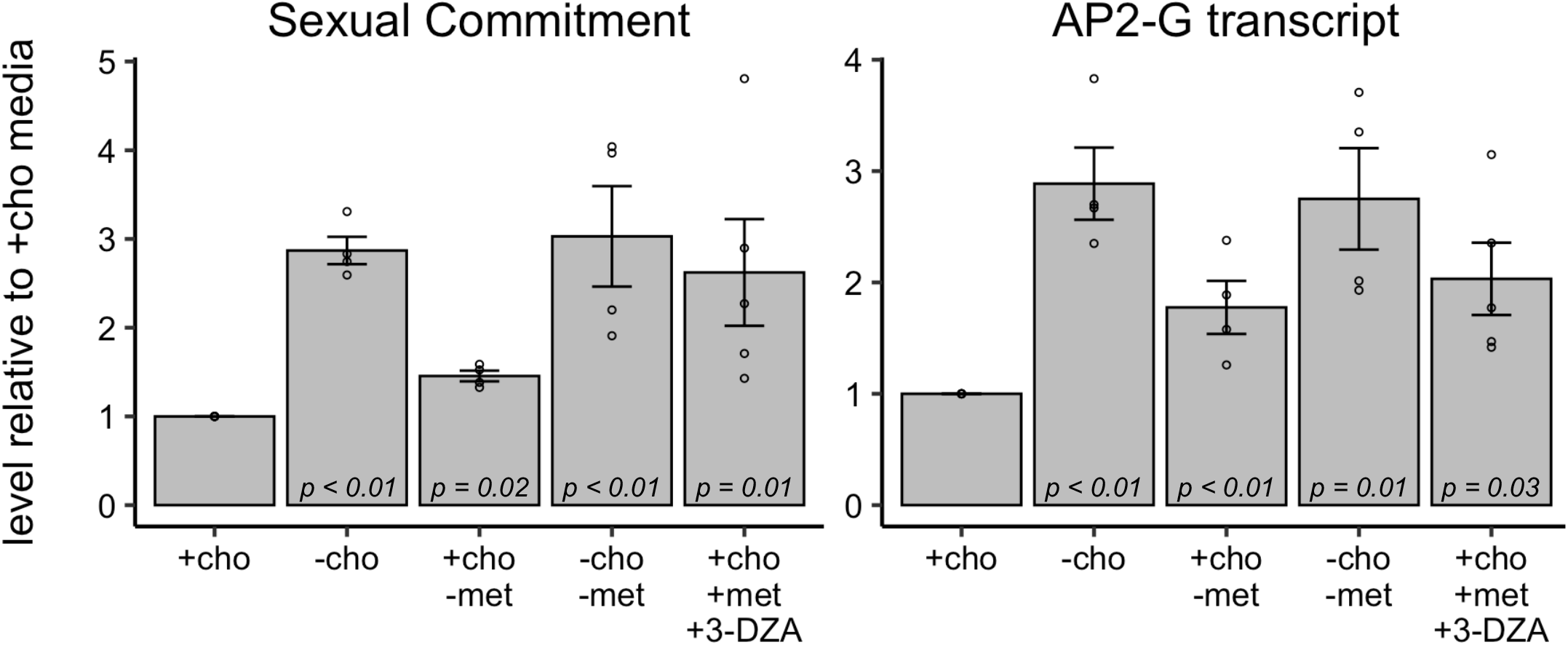
Schizont AP2-G transcript abundances closely track sexual commitment under various nutrient conditions. Bars indicate the mean sexual commitment (left) and AP2-G transcript abundance (right) in schizonts relative to conditions of abundant choline and methionine (+cho) when parasites where exposed to different growth media during the commitment cycle. Error bars and p-values indicate the standard error of the mean and the significance of the mean difference relative to those under conditions of abundant choline and methionine (+cho+met), respectively. n=4-5.

**Extended Data Figure 4:**
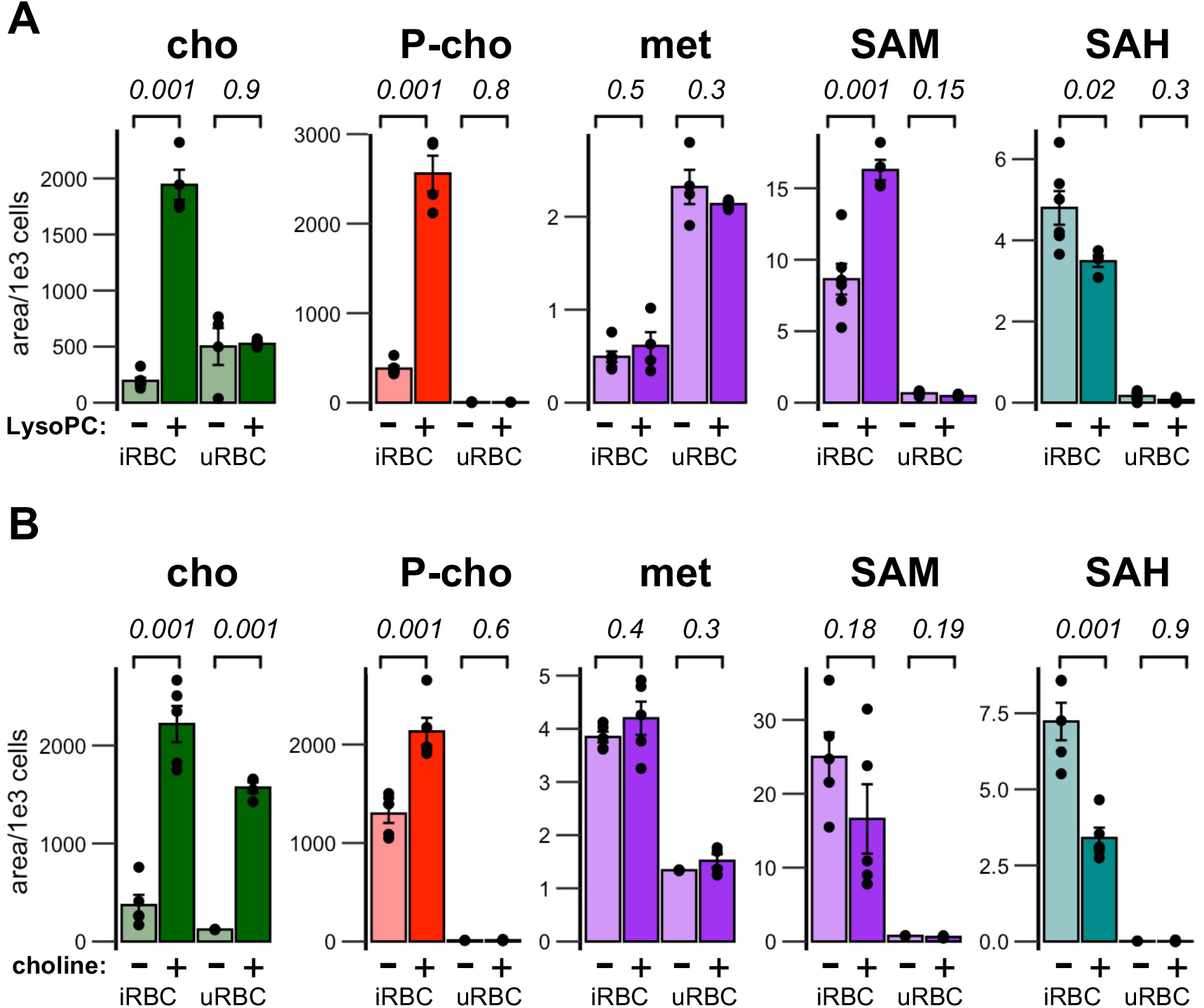
Changes in SAM/SAH metabolism are specific to parasite metabolism. LCMS quantification of indicted metabolites. Infected and uninfected cultures were cultured in the presence or absence of 20 μM LysoPC or 420μM choline for ~36 hpi during the commitment cycle. Infected (iRBC) and uninfected (uRBC) erythrocytes were then extracted, and metabolite abundances were quantified by LCMS. Bar graphs show the mean intracellular metabolite abundances per thousand cells ± s.e.m (n=3-5). Italicized numbers are p-values based on two-sided paired t-tests.

**Extended Data Figure 5:**
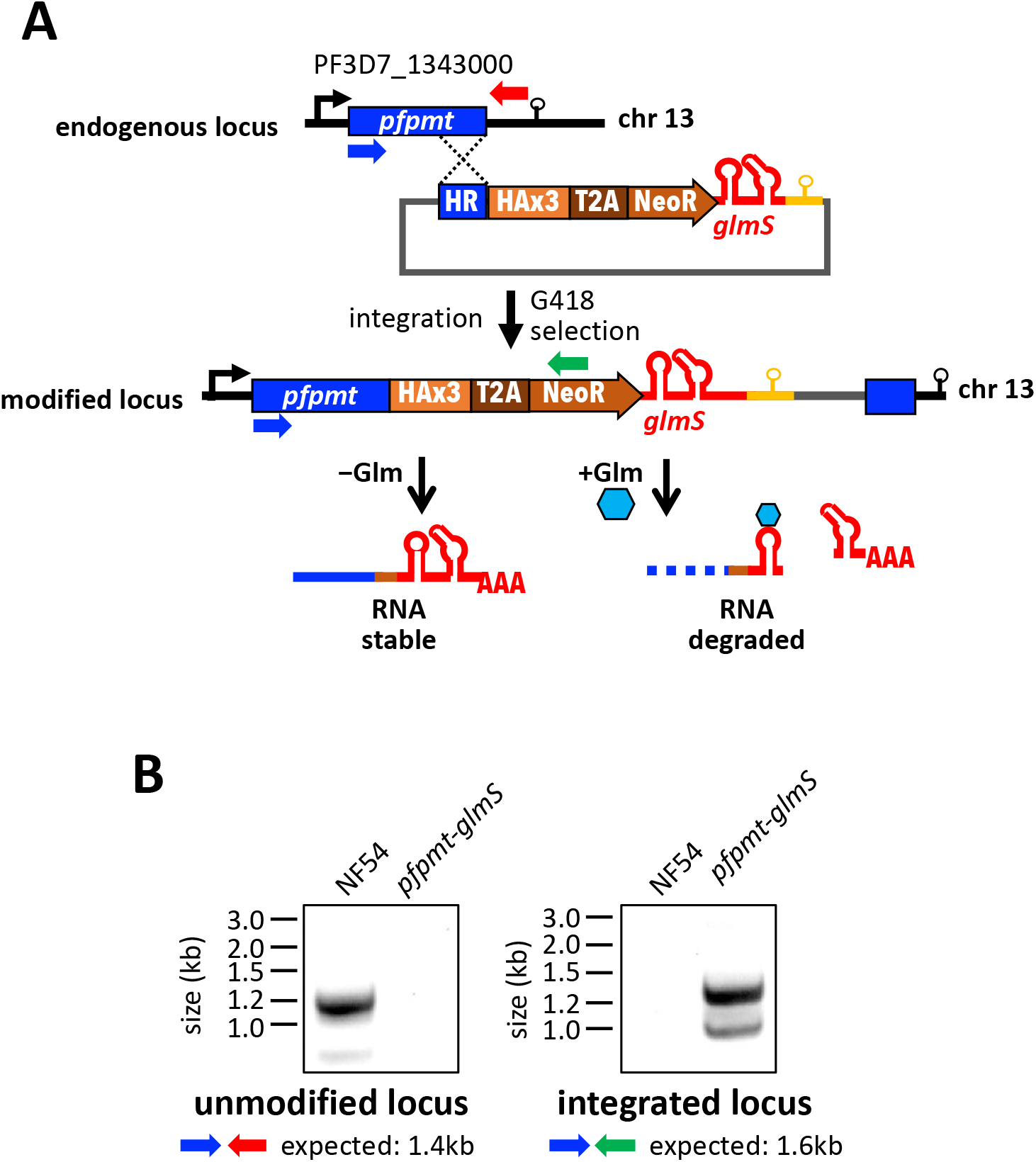
Validation of *PMT-glmS* knockdown parasite line. **(A)** Generation of *PMT-glmS* knockdown parasites by selection-linked integration. **(B)** Validation PCR demonstrating tagging of the endogenous *PMT* locus.

**Extended Data Figure 6:**
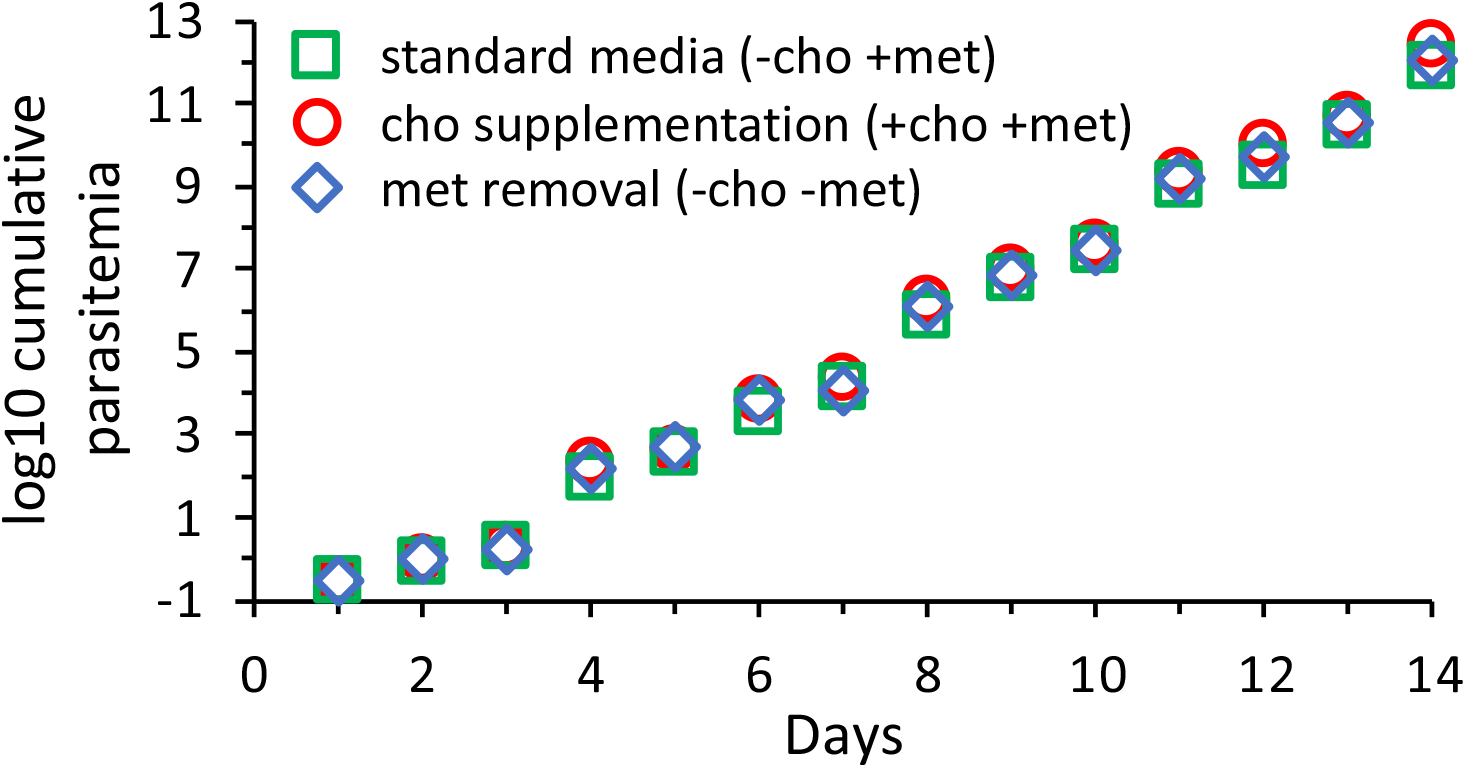
Removal of methionine (blue diamond) or supplementation with choline (red circles) had no observable effect on growth of NF54 compared to growth in standard malaria medium (green squares).

**Extended Data Figure 7:**
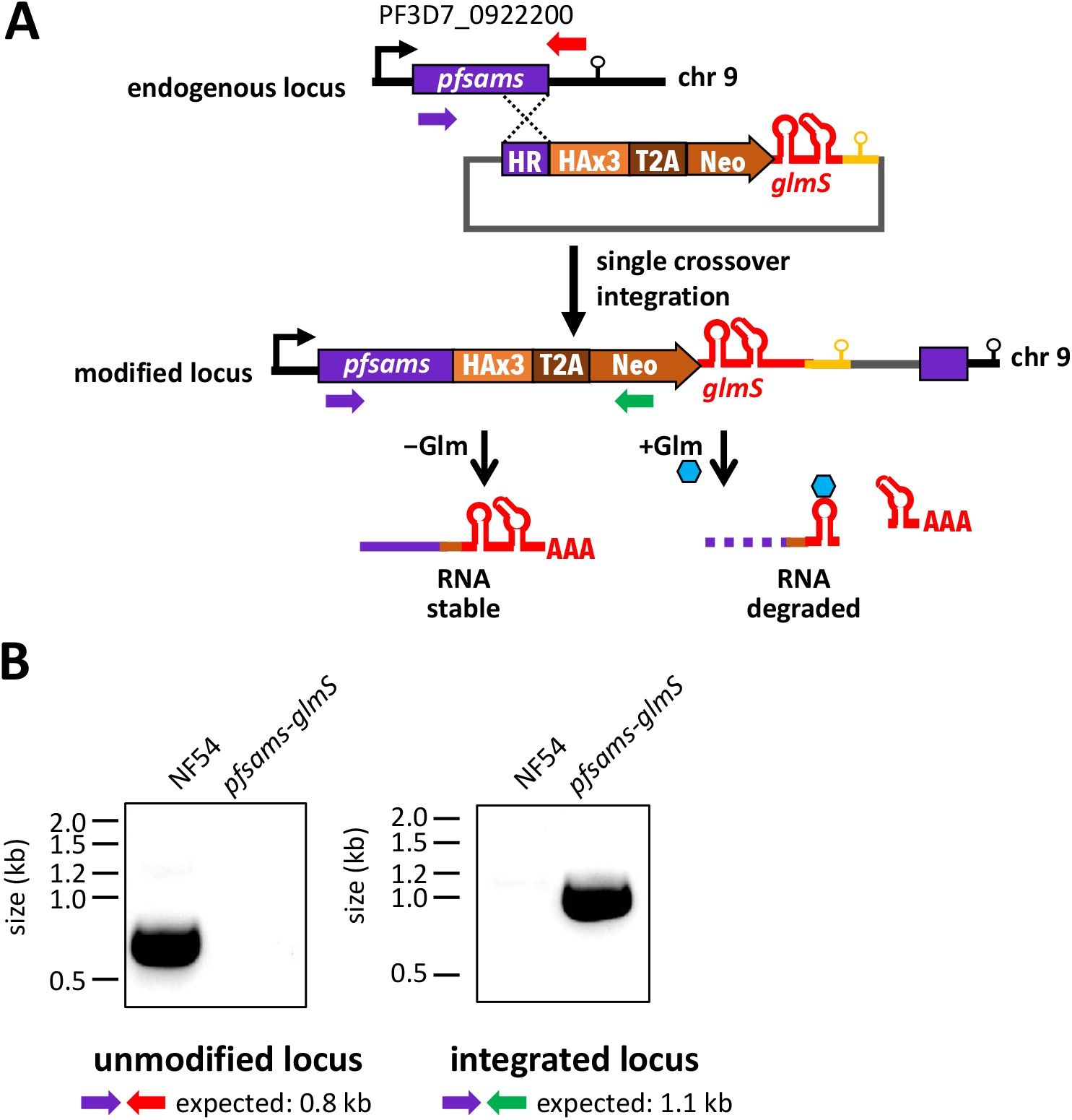
Validation of *pfsams-glmS* knockdown parasite line. **(A)** Generation of *pfsams-glmS* knockdown parasites by selection-linked integration. **(B)** PCR Validation demonstrating tagging of the endogenous *pfsams* locus.

**Extended Data Figure 8:**
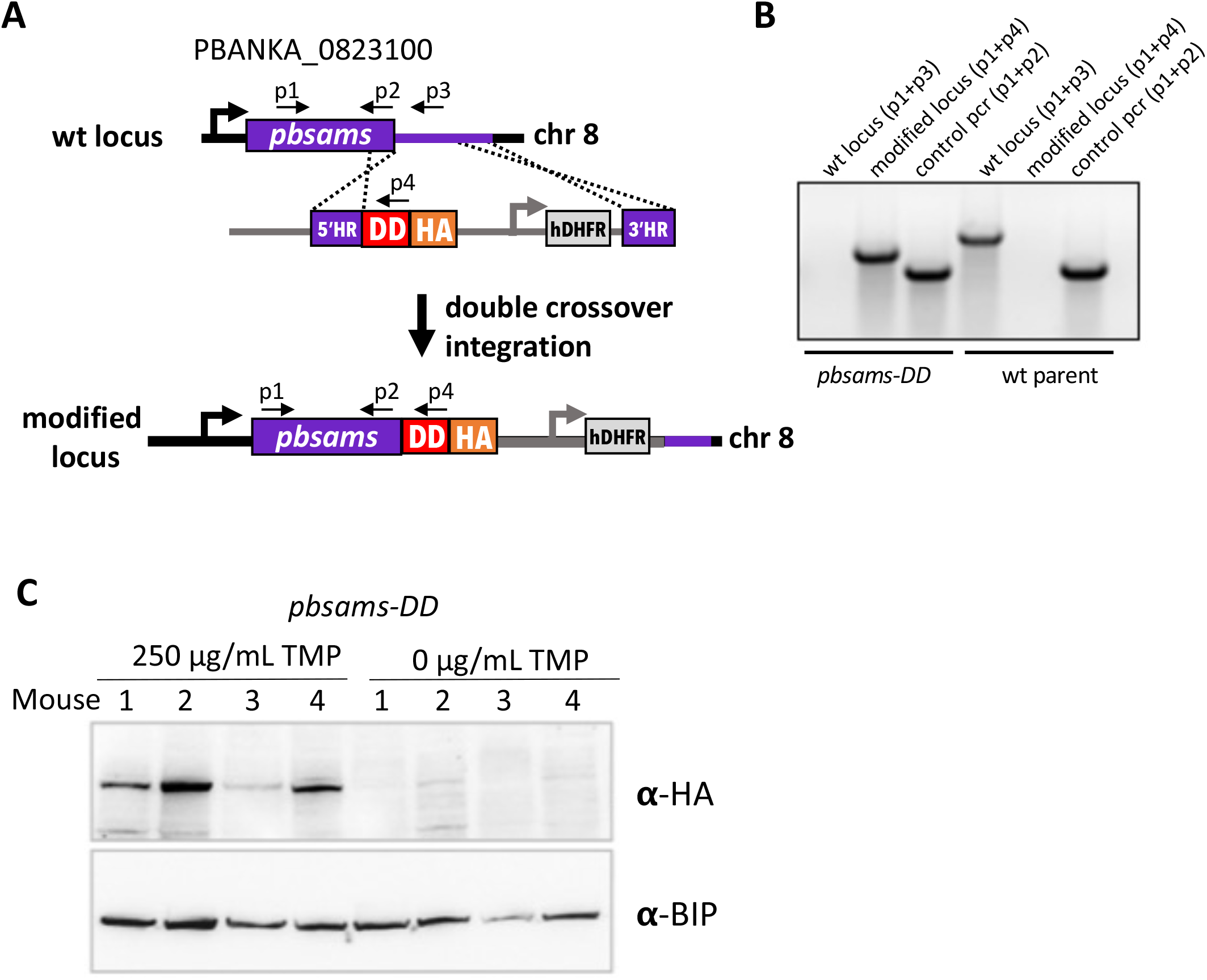
Validation of *pbsams-DD* knockdown parasite line. **(A)** The endogenous *pbsams* locus in the *P. berghei* ANKA strain background was modified by homologous integration to add the ecDHFR destabilization domain (DD) and hemagglutinin epitope tag (HA) at the 3’ end of the *pbsams* coding sequence. Simultaneous integration of a hDHFR expression cassette allows for selection of integrants. **(B)** PCR validation of successful tagging in *Pb*SAMS-DD-HA parasites. **(C)** Successful knockdown of *Pb*SAMS upon removal of trimethoprim (TMP) from the drinking water in mice infected with *pbsams-DD* parasites. Parasite lysates were assayed for the abundance of *Pb*SAMS-DD with antibodies against the HA epitope tag and *Pb*BIP, which served as a loading control and was used for normalization.

**Extended Data Figure 9:**
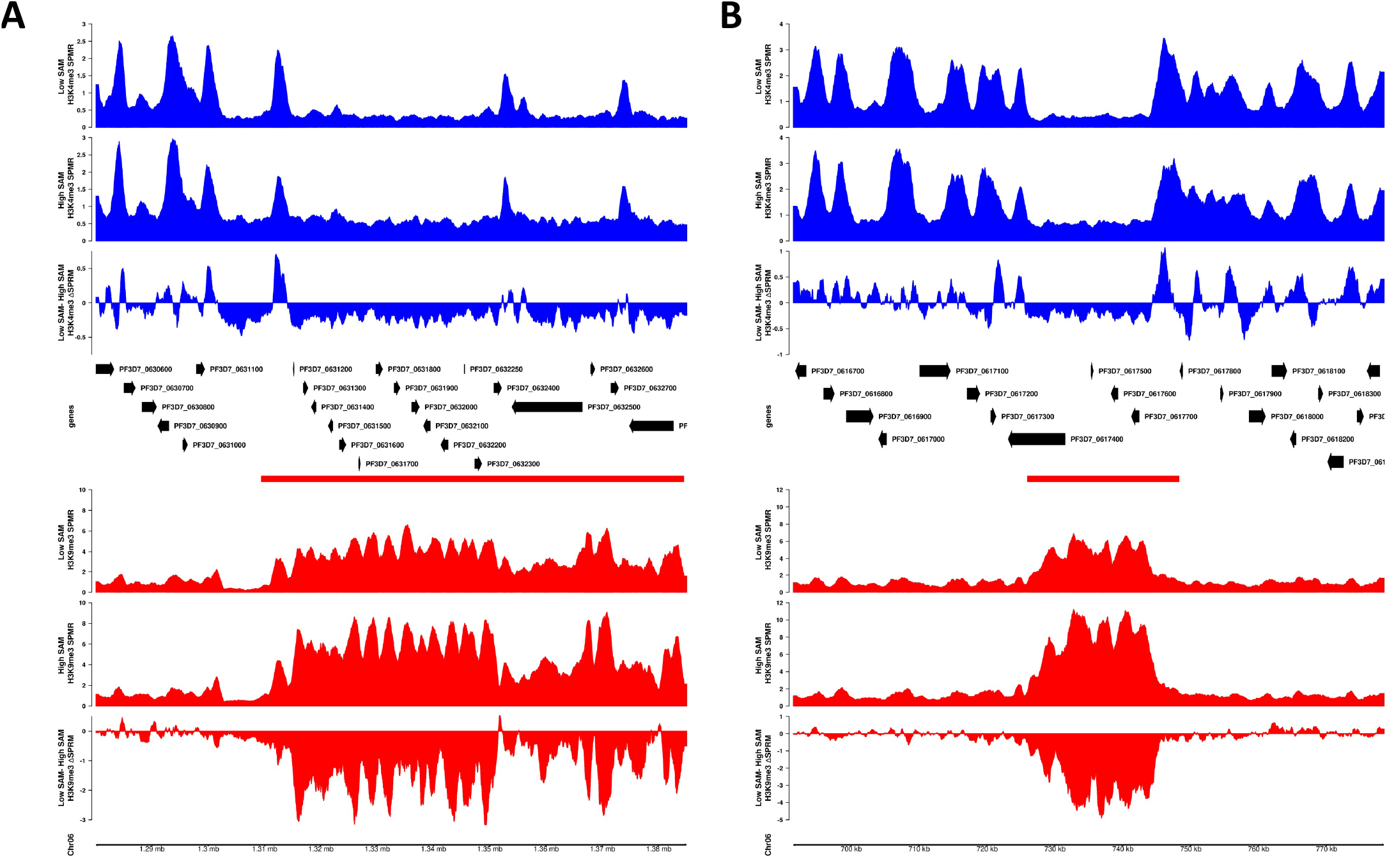
Coverage comparisons of H3K4me3 and H3K9me3 at two representative chromosome 6 loci under Low vs. High SAM conditions. Coverage of H3K4me3 (blue) and H3K9me3 (red) at representative regions on chromosome 6 that include euchromatin and either subtelomeric heterochromatin **(A)** or a heterochromatin island **(B)** under Low SAM (top track of each color) and High SAM conditions (middle track of each color) and the relative difference in coverage (third track of each color). Heterochromatin regions are marked with a red bar. Coverage was normalized as signal per million reads (SPRM) using macs2.

**Extended Data Figure 10:**
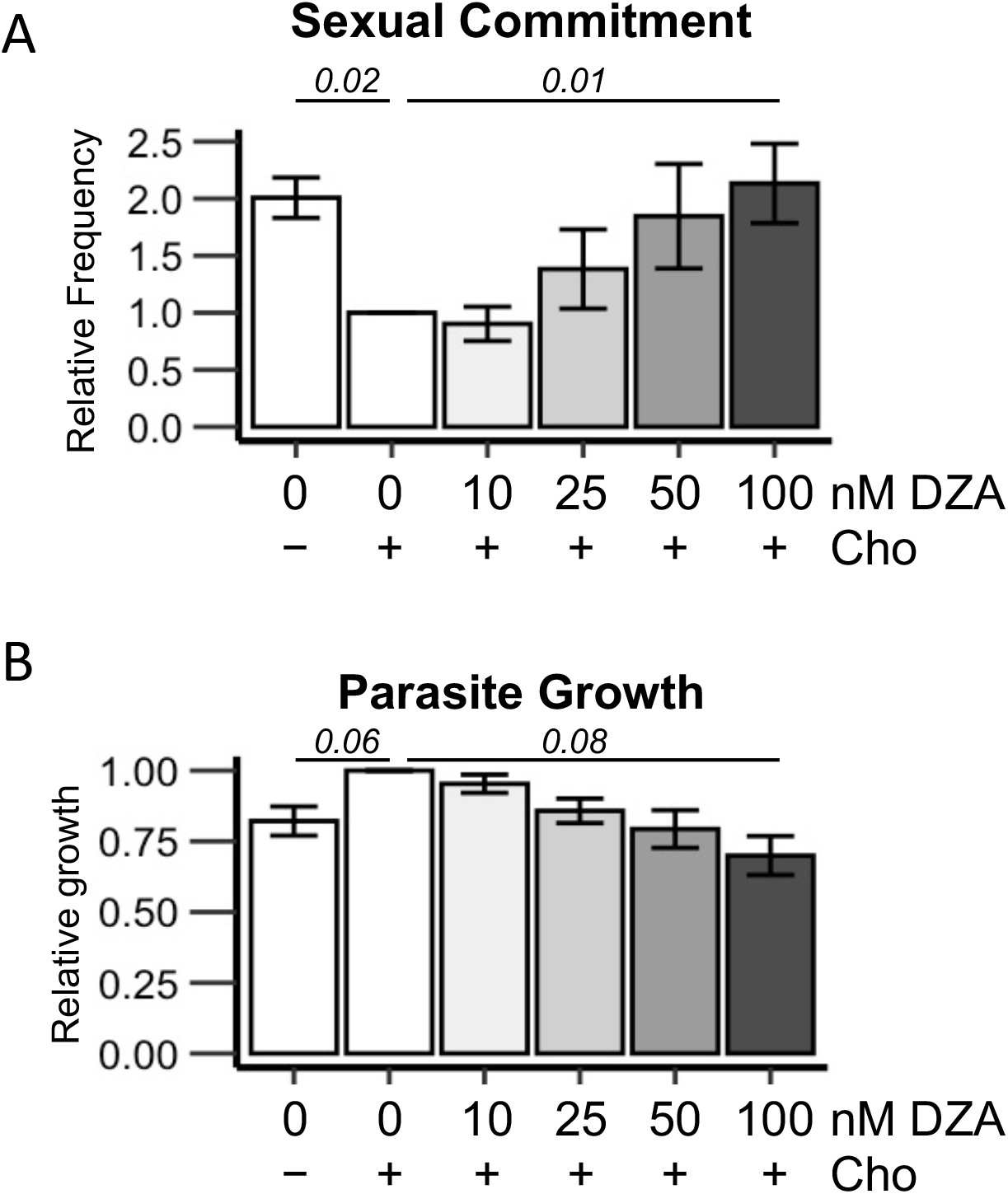
Dose-response of parasite sexual commitment (A) and growth (B) to 3-DZA. Italicized number is the p-value based on a two-sided t-tests for the +/- choline comparison and ANOVA for the DZA dose response (n=4).

**Extended Data Figure 11:**
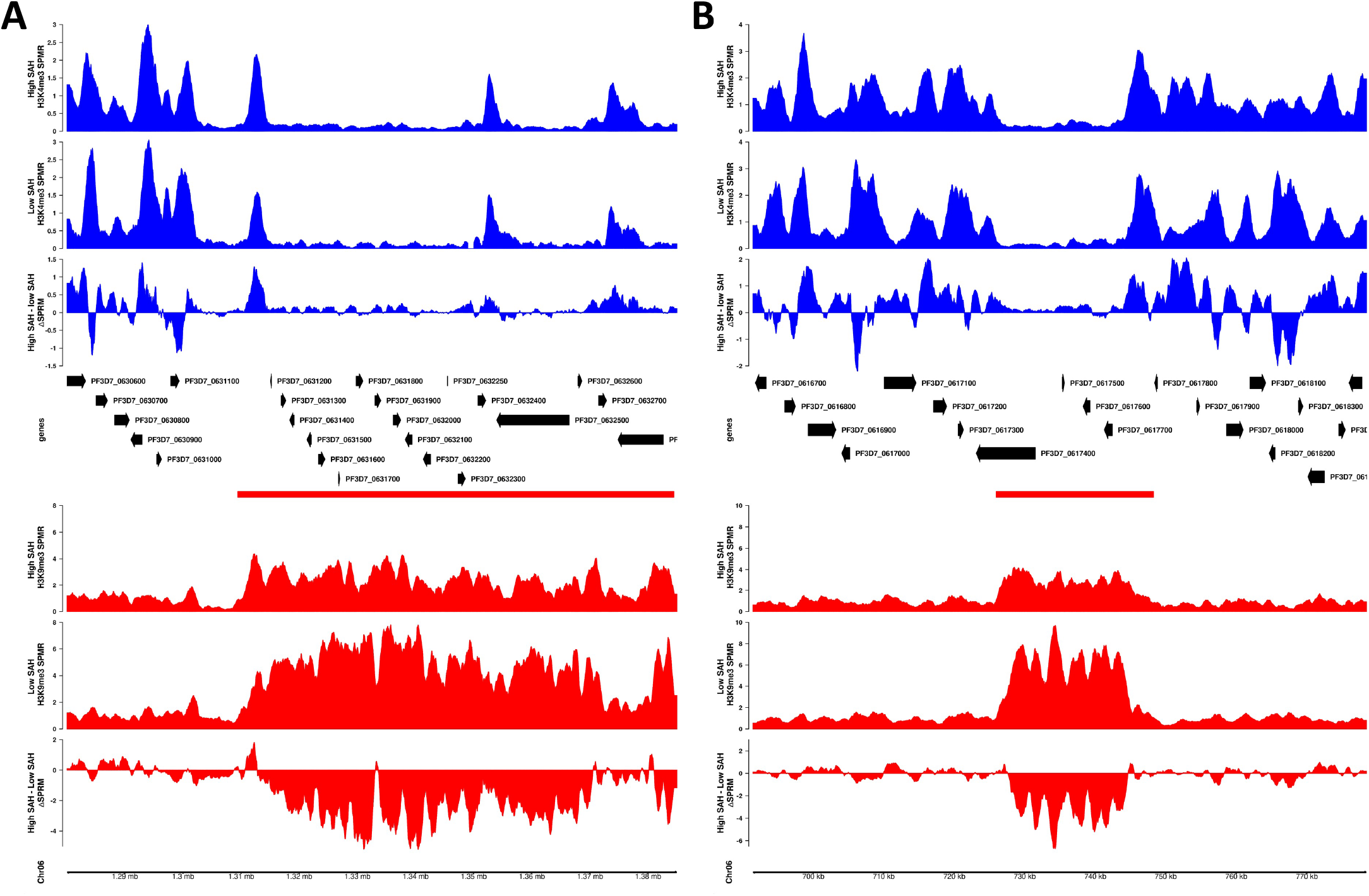
Coverage comparisons of H3K4me3 and H3K9me3 at two representative chromosome 6 loci under High vs. Low SAH conditions. Coverage of H3K4me3 (blue) and H3K9me3 (red) at representative regions on chromosome 6 that include euchromatin and either subtelomeric heterochromatin **(A)** or a heterochromatin island **(B)** under High SAH (top track of each color) and Low SAH conditions (middle track of each color) and the relative difference in coverage (third track of each color). Heterochromatin regions are marked with a red bar. Coverage was normalized as signal per million reads (SPRM) using macs2.

**Extended Data Table 1:** Compositions of cell culture media used in this study.

| Label | Differences to standard malaria complete media |
| --- | --- |
| - cho | 0 µM choline |
| + cho | 420 µM Choline |
| - LysoPC | 30 µM Palmitate, 30 µM oleate, 1.25 µM LysoPC, 4 g/L Fatty Acid Free BSA, no Albumax II, |
| + LysoPC | 30 µM Palmitate, 30 µM oleate, 40 µM LysoPC, 4 g/L Fatty Acid Free BSA, no Albumax II, |
| - Glm | none |
| + Glm | 2.5mM glucosamine |
| - met | 0 µM methionine |
| + met | none (100 µM methionine) |
| - 3DZA | 420 µM Choline |
| + 3DZA | 420 µM Choline, 100nM 3-DZA |

**Extended Data Table 2:**
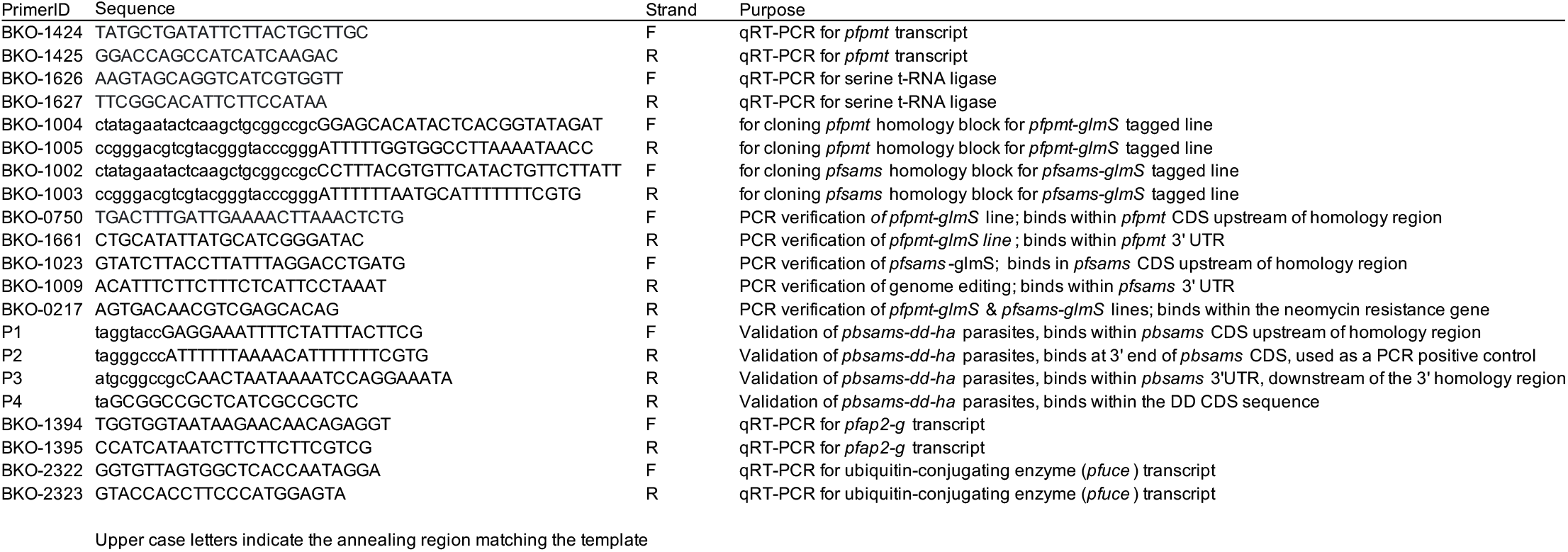
Primers used in this study

